# Coronavirus papain-like protease antagonizes innate immunity by cleaving Importin α1 to disrupt nuclear transport

**DOI:** 10.64898/2026.07.17.739146

**Authors:** Jiehuan Wang, Wenxiang Xue, Yingjie Sun, Lei Tan, Cuiping Song, Xusheng Qiu, Chan Ding, Ying Liao

## Abstract

The Importin α family, as key mediators of nucleocytoplasmic transport, represents a common target for viral immune evasion. However, whether coronaviruses directly manipulate Importin α to disrupt nuclear trafficking and suppress antiviral immunity has remained unclear. In this study, we identify a previously unrecognized mechanism by which coronaviruses from all four genera subvert host innate immunity through the proteolytic inactivation of Importin α1, a key mediator of nuclear import of cargo proteins. We demonstrate that the membrane-associated papain-like protease (PLpro-TM) directly cleaves Importin α1 at specific glycine residues, G129 for PEDV and IBV PLpro, and G119 for MHV and PDCoV PLpro, thereby disrupting its nuclear import function. This cleavage impairs the nuclear translocation of multiple transcription factors (IRF3, STAT1, STAT2, and p65) and suppresses the expression of downstream antiviral genes, including IFN-β and IFN-stimulated genes (ISGs). Importantly, cleavage-resistant mutants of Importin α1 (G129A or G119A) restore nuclear import capability and rescue IFN-β signaling. Consequently, they exert a more potent inhibitory effect on viral replication than the wild-type Importin α1, fulfilling an antiviral role. Our work establishes PLpro-TM-mediated cleavage of Importin α1 as a conserved immune evasion strategy across coronaviruses and highlights this interaction as a potential target for broad-spectrum antiviral intervention.

**Author summary:** The nuclear transport of transcription factors is a critical checkpoint for the initiation of innate antiviral immunity. Here, we identify the PLpro-TM protein as a pan-coronavirus antagonist of nucleocytoplasmic trafficking and innate immune response. We demonstrate that PLpro-TM from four distinct coronavirus genera directly cleaves Importin α1 at specific glycine residues, thereby disabling its ability to mediate the nuclear import of key transcription factors and subsequent transcription of anti-viral genes. This work reveals a previously unrecognized, evolutionarily conserved immune evasion strategy shared across α, β, γ, and δ coronaviruses. By uncovering the proteolytic inactivation of Importin α1 targeting by PLpro-TM, our findings not only resolve a long-standing question about how coronaviruses disrupt nuclear trafficking, but also establish PLpro-TM and Importin α1 as promising targets for the development of broad-spectrum antiviral therapeutics against current and emerging coronaviruses.

## Introduction

The innate immune system serves as the first line of defense against virus infection. Upon detection of pathogen-associated molecular patterns, pattern recognition receptors activates downstream signaling cascades that culminate in the nuclear translocation of transcription factors, notably IRF3, STAT1/STAT2/IRF9, and NF-κB, to induce type I/III interferons, interferon-stimulated genes (ISGs), and pro- or anti-inflammatory cytokines [1–5]. The classical nuclear import pathway, mediated by the Importin α/β1 heterodimer, is central to this process: Importin α recognizes the nuclear localization signal (NLS) of cargo proteins, while Importin β1 facilitates their translocation through the nuclear pore complex [6–9]. Given its essential role in antiviral immunity, the nuclear transport machinery has emerged as a frequent target of viral antagonism [10–12].

There are seven human Importin α isoforms, each of which exhibits distinct binding affinities for specific NLSs, thereby conferring cargo selectivity and transport specificity [12, 13]. For example, upon activation, the transcription factor IRF3 drives the expression of type I and III interferon, initiating the first line of host defense [14]. The nuclear translocation of activated IRF3 is mediated by the classical Importin α/β1 pathway, involving Importins α1, α3, and α4 [15–17]. Once secreted, type I interferons bind to cell-surface receptors and activate the JAK/STAT signaling pathway [18]. This cascade ultimately leads to the nuclear translocation of phosphorylated STAT1-STAT2 heterodimers, which drive the expression of a broad spectrum of ISGs [19]. Notably, the nuclear import of the STAT1 complex is specifically mediated by Importins α1, α5, α6, and α7 [20–22]. In contrast, STAT2 depends on the formation of the ISGF3 complex with STAT1 and IRF9 for efficient nuclear translocation, which typically occurs via Importin α5 [22, 23]. Similarly, activation and nuclear translocation of NF-κB (p50/p65) drive the transcription of antiviral and inflammatory cytokines [24], a process that relies on Importins α1, α3, and α4, together with Importin β1, for nuclear entry [16, 17, 25–27]. Collectively, Importins α and β represent critical host factors that viruses can exploit to impede the nuclear translocation of essential transcription factors and thereby antagonize the host antiviral response. Among the various Importin α isoforms, Importin α1 occupies a particularly strategic position in antiviral signaling. As noted above, it serves as a shared nuclear transporter for multiple key transcription factors, including IRF3, STAT1, and the p65 subunit of NF-κB [12, 13]. Consequently, disruption of Importin α1 function would be expected to broadly impair innate immune responses, a vulnerability that several viruses have evolved to exploit.

Coronaviruses are enveloped, positive-sense RNA viruses that pose significant threats to both human and animal health, as exemplified by severe acute respiratory syndrome coronavirus 2 (SARS-CoV-2), middle east respiratory syndrome coronavirus (MERS-CoV), and, in the veterinary context, porcine epidemic diarrhea virus (PEDV) and infectious bronchitis virus (IBV) [28–31]. The α-coronavirus PEDV is a major pathogen in swine, causing acute, severe enteric disease characterized by watery diarrhea, vomiting, dehydration, and high mortality in neonatal piglets, resulting in substantial economic losses to the global swine industry [29, 32–34]. Similarly, γ-coronavirus IBV is a highly contagious that poses a major threat to the poultry industry. This virus primarily causes respiratory disease in chickens but can also infect the renal, reproductive, and digestive tracts, leading to reduced egg production and significant economic losses in the poultry industry worldwide [35–37]. This virus exhibits extensive genetic diversity, giving rise to numerous serotypes and variants that continuously challenge current vaccination strategies [38–40]. Porcine δ-coronavirus (PDCoV) is an emerging enteric pathogen that poses a growing threat to the global swine industry. It primarily causes acute diarrhea, vomiting, and intestinal damage in piglets, leading to substantial economic losses [41–43].

To establish successful infection, coronaviruses have evolved diverse strategies to subvert the host nuclear transport machinery, blocking the nuclear entry of essential transcription factors and thereby evading innate immunity [6, 44]. For instance, the β-coronavirus SARS-CoV-2 accessary protein ORF6 blocks STAT1 import by binding Rae1 and also interacts with Importin α1 to inhibit Importin α/β1-mediated IRF3 and STAT1 nuclear translocation [45–47]; MERS-CoV accessary protein 4b competitively binds Importin α3 to prevent the p65 nuclear import [48]; PEDV nsp7 specifically blocks the interaction between Importin α5 and STAT1 to suppress STAT1 nuclear entry [49]; and PDCoV nucleocapsid (N) protein promotes Importin α1 degradation to suppress STAT1 nuclear import [20]. More broadly, our recent report revealed that pan-coronavirus N protein disrupts the nucleocytoplasmic transport by inducing the phosphorylation of nucleoporin 62, thereby suppressing the innate immune response [50]. Despite this growing list of examples, the mechanistic repertoire employed by coronaviruses to disrupt nuclear transport remains incompletely defined, and whether additional viral proteins directly target Importin α family members has not been explored.

The papain-like protease (PLpro) is a multifunctional enzyme conserved across the *Coronaviridae* family, possessing both deubiquitinating and proteolytic activities that are essential for viral polyprotein processing and immune evasion [51, 52]. PLpro has been shown to antagonize type I interferon responses by removing ubiquitin and ISG15 modifications from key signaling molecules [53–55]. However, whether PLpro exerts additional functions, particularly through direct cleavage of host proteins involved in nuclear transport, has remained unknown. In this study, we identify the membrane-anchored form of coronavirus PLpro (PLpro-TM) as a viral effector that directly cleaves Importin α1 to disrupt nucleocytoplasmic trafficking and suppress innate immunity. We demonstrate that PLpro-TM reduces Importin α1 protein levels and alters its subcellular distribution, thereby blocking the nuclear import of IRF3, STAT1/STAT2, and p65, and attenuating antiviral gene expression. Mechanistically, PLpro-TM cleaves Importin α1 at specific glycine residues, G129 in the cases of PEDV and IBV, and G119 for MHV and PDCoV, revealing both conserved and virus genus-specific cleavage preferences. The conservation of this cleavage across α-, β-, and γ-coronaviruses suggests that PLpro-mediated Importin α1 cleavage represents a broad-spectrum viral strategy to subvert host defenses. Our findings not only expand the mechanistic understanding of coronavirus-host interactions but also identify PLpro-TM and Importin α1 as potential targets for the development of pan-coronavirus antiviral therapeutics.

## Results

### PEDV infection reduces Importin α1 and α4 expression and increase their cytoplasmic accumulation

To investigate whether coronavirus infection affects Importin-mediated nuclear transport, Vero cells were infected with the PEDV HLJBY strain, and the expression levels of multiple Importin α isoforms (α1, α3, α4, α5, α6, and α7) were examined via Western blotting. As shown in Fig 1A, PEDV infection induced a progressive and marked decrease in Importin α1 levels, alongside a moderate reduction in Importin α4, from 12 and 24 hours post-infection (h.p.i.). Notably, an additional smaller band below the full-length Importin α4 appeared at 24 h.p.i., suggesting possible cleavage or degradation. In contrast, the protein levels of Importin α3, α5, α6, and α7 remained relatively stable throughout the infection time course. To further assess the impact of PEDV infection on the subcellular localization of Importin α1 and Importin α4, immunofluorescence staining was performed on PEDV-infected Vero cells. In mock-infected cells, both Importin α1 and Importin α4 were predominantly localized in the nucleus. However, upon PEDV infection, a clear cytoplasmic accumulation of both proteins was observed from 6 to 18 h.p.i. (Fig 1B). Collectively, these results demonstrate that PEDV infection not only downregulates the expression of Importin α1 and Importin α4 but also disrupts their normal nucleocytoplasmic shuttling, potentially impairing host cellular transport processes.

**Fig 1.**
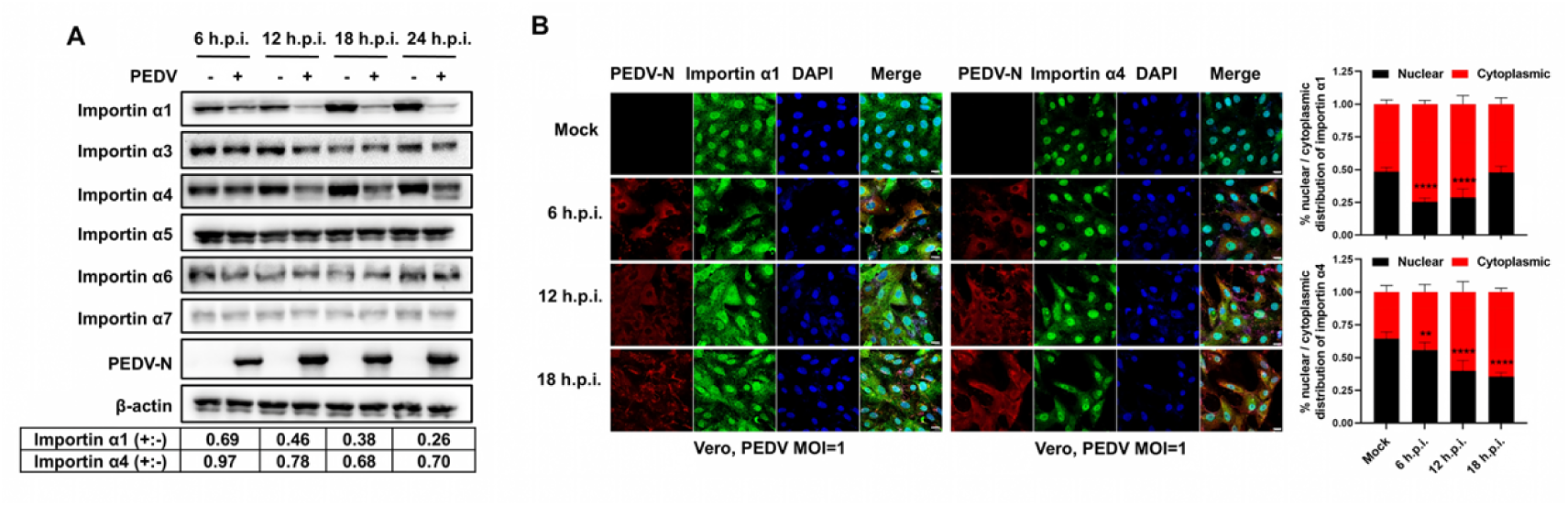
PEDV regulates Importin α1 and Importin α4 expression and subcellular localization. (A) Vero cells were infected with PEDV (MOI = 1) and harvested at 6, 12, 18, 24 h.p.i.. Cell lysates were subjected to Western blot analysis to detect Importin α1, α3, α4, α5, α6, and α7. β-actin served as a loading control. The band intensities for Importin α1 and Importin α4 were measured using ImageJ, normalized to β-actin, and are presented as the ratio of virus-infected to mock-infected samples. (B) Vero cells were infected with PEDV and harvested at 6, 12, 18 h.p.i.. Immunofluorescence analysis was performed to detected viral N protein (red) and Importin α1 or Importin α4 (green). Nuclei were counterstained with DAPI (blue). Representative images were shown. Scale bar, 10 μm. 10 cells were selected from three distinct fields of view in mock-infected or PEDV-infected cells, and the nuclear and cytoplasmic Importin α1 or Importin α4 fluorescence signals were quantified using ImageJ. The bar graphs depict the mean signal intensity in the nucleus (black bars) and cytoplasm (red bars), with error bars representing the standard deviation (SD). *P* values were determined by Student’s t-test: **, *P* < 0.01; ****, *P* < 0.0001.

### Coronavirus PLpro-TM specifically targets Importin α1 for downregulation and cytoplasmic accumulation

To identify the viral proteins responsible for targeting Importin α1, we generated expression plasmids encoding individual PEDV proteins, including nsp1–nsp16, S, N, E, M, and ORF3, each with an N-terminal Flag tag. Given that nsp3 is the largest viral protein and its full-length cloning is technically challenging, we cloned only the PLpro-TM region, which is located at the C-terminus of nsp3 and comprises the PLpro2 domain, the ectodomain (3Ecto), and two transmembrane regions (TM1 and TM2), to evaluate its effect on Importin α1 expression. To ensure efficient transfection and protein expression, HEK-293T cells were co-transfected with plasmids expressing each viral protein alongside Importin α1 (N-terminally HA-tagged and C-terminally Myc-tagged). The expression levels of both Importin α1 and the viral proteins were subsequently assessed. As shown in Fig 2A, most viral proteins were successfully expressed, with the exception of nsp7, nsp9, nsp12 and S. Notably, nsp1, PLpro-TM, nsp9, nsp13, and nsp15 markedly reduced Importin α1 expression, with PLpro-TM exerting the most pronounced suppressive effect. Given that nsp1 and nsp15 are known global translation suppressors [56–58], we focused our subsequent mechanistic analysis on PLpro-TM, which demonstrated potent effect on Importin α1.

To determine whether Importin α1 targeting by PLpro-TM is conserved across four coronavirus genera, we extended our analysis to include PLpro-TM from representatives of three additional genera: β-coronavirus Murine Hepatitis Virus (MHV), γ-coronavirus IBV, and δ-coronavirus PDCoV. MHV is a prototypic β-coronavirus that has long served as an indispensable experimental model for studying coronavirus biology, pathogenesis, and host immune responses [59, 60]. Each construct encompasses the region spanning from the PLpro2 domain through TM1 and TM2 (Fig 2B), with a Flag tag fused to its N-terminus. When co-expressed with HA-Importin α1-Myc in HEK-293T cells, PLpro-TM from all four coronaviruses (PEDV, MHV, IBV, PDCoV) significantly reduced the exogenous expressed Importin α1 levels (Fig 2C). Furthermore, prolonged exposure of the immunoblot revealed an additional ∼45 kDa band corresponding to Importin α1 in cells co-expressing PLpro-TM, suggesting the generation of a cleavage product. This ∼45 kDa fragment was detected by the C-terminal tag antibody (anti-Myc), confirming that it represents the C-terminal portion of Importin α1. We next tested the endogenous Importin α1 reduction by PLpro-TM from PEDV and IBV. As shown in Fig 2D, increasing expression of PLpro-TM caused a progressive decrease in endogenous Importin α1 levels. These results demonstrate that the specific proteolytic cleavage of Importin α1 by PLpro-TM is a conserved mechanism across coronaviruses. We also tested the effect of PLpro-TM on Importin α4 expression level. Only PEDV PLpro-TM reduced the levels of co-expressed Importin α4, whereas MHV, IBV, and PDCoV PLpro-TM had no detectable effect (S1A Fig).

To determine whether PLpro-TM alters endogenous Importin α1 localization, we performed immunofluorescence analysis. In vector PXJ40 transfected cells or PLpro-TM absent cells, endogenous Importin α1 accumulated in the nucleus; however, expression of PLpro-TM from PEDV, MHV, IBV, and PDCoV consistently caused Importin α1 to retain in the cytoplasm (Fig 2E). Interestingly, we observed significant co-localization between Importin α1 and PLpro-TM, indicative of an interaction (Fig 2E). Examination of Importin α4 localization revealed its robust nuclear import regardless of PLpro-TM expression (S1B Fig), consistent with the minimal impact of PLpro-TM on Importin α4 protein levels (S1A Fig). Collectively, these data support a mechanism whereby coronavirus PLpro-TM specifically targets and cleaves Importin α1, consequently blocking its nuclear translocation.

**Fig 2.**
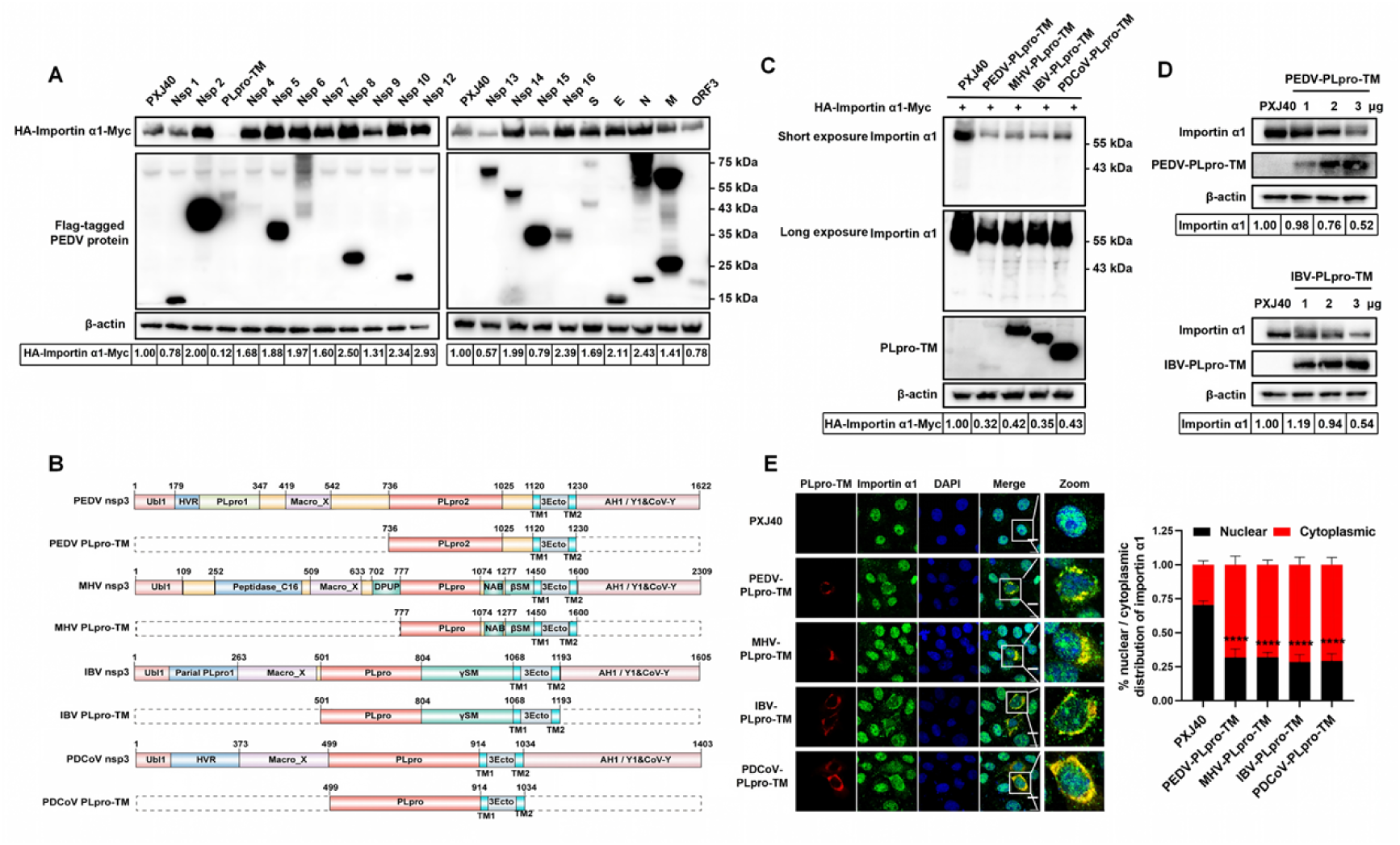
Coronavirus PLpro-TM targets Importin α1 for cleavage, reducing its expression and causing cytoplasmic accumulation. (A) Screen for viral proteins that alter Importin α1 expression. HEK-293T cells were co-transfected with plasmids encoding Importin α1 (with HA tag and Myc tag) and Flag-tagged viral proteins from PEDV. Lysates were analyzed by Western blot with antibodies against Myc, Flag, and β-actin. (B) Schematic of coronavirus nsp3, highlighting the domain architecture of the PLpro-TM constructs from PEDV, MHV, IBV, and PDCoV. Nsp3 key domains: Ubiquitin-like 1 (Ubl1), hypervariable region (HVR), macrodomain (Macro), PLpro, nucleic acid-binding domain, beta-coronavirus-specific marker (βSM), gamma-coronavirus-specific marker (γSM), TM1, TM2, nsp3 ectodomain (3Ecto), Y1, and CoV-Y. (C) PLpro-TM from pan-coronaviruses downregulates Importin α1. HEK-293T cells were co-transfected with HA-Importin α1-Myc and Flag-PLpro-TM from PEDV, MHV, IBV, and PDCoV. Cell lysates were analyzed with anti-Flag, anti-Myc, or anti-β-actin antibodies. (D) PLpro-TM reduces the level of endogenous Importin α1. HEK-293T cells were transfected with increasing amounts of PEDV or IBV PLpro-TM plasmids. After 24 h, lysates were analyzed by Western blot for detecting endogenous Importin α1, PLpro-TM, and β-actin. The Importin α1 protein band densities in panel A, C, D were quantified with ImageJ, normalized to β-actin, and are presented as ratios relative to the PXJ40 group. (E) PLpro-TM mediates the cytoplasmic retention of endogenous Importin α1. Vero cells were transfected with Flag-tagged PLpro-TM constructs from PEDV, MHV, IBV, or PDCoV. At 24 hours post-transfection (h.p.t.), cells were fixed and stained for the Flag-PLpro-TM (red) and endogenous Importin α1 (green). Nuclei were counterstained with DAPI (blue). Representative images from three independent experiments are shown. Scale bar, 10 μm. The nuclear and cytoplasmic Importin α1 fluorescence signals were quantified using ImageJ in 10 cells selected from distinct fields of view, comparing control (PXJ40) and PLpro-TM-expressing cells. The bar graphs depict the mean signal intensity in the nucleus (black bars) and cytoplasm (red bars), with error bars representing the SD. *P* values were determined by Student’s t-test: ****, *P* < 0.0001.

**S1 Fig.**
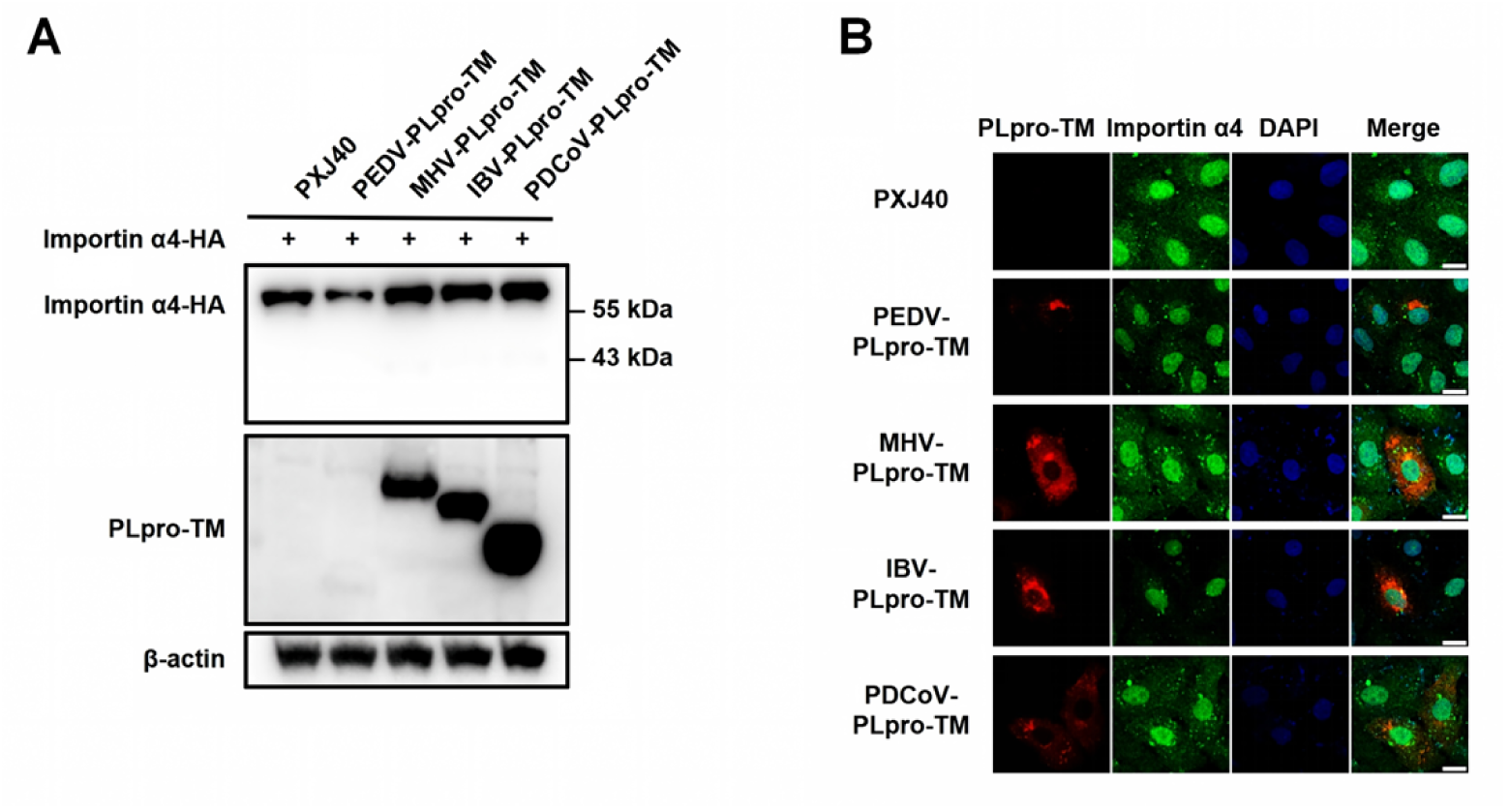
Coronavirus PLpro-TM does not alter the expression level and subcellular localization of Importin α4. (A) PLpro-TM has minimal effect on Importin α4 expression. HEK-293T cells were co-transfected with HA-tagged Importin α4 and Flag-tagged PLpro-TM from PEDV, MHV, IBV, PDCoV. Cell lysates were analyzed by Western blot for HA-Importin α4, Flag-PLpro-TM, and β-actin. (B) Importin α4 nuclear entry is not interfered by PLpro-TM. Vero cells expressing Flag-tagged PLpro-TM from the indicated coronaviruses were subjected to immunostaining for endogenous Importin α4 (green) and Flag-PLpro-TM (red) at 24 h.p.t.. Nuclei were counterstained with DAPI (blue). Scale bars: 10 µm.

### Interaction between coronavirus PLpro-TM and Importin α1

The observed cleavage of Importin α1 in PLpro-TM-expressing cells led us to hypothesize a direct interaction between these two proteins. To test this, we performed co-immunoprecipitation using exogenous expressed Flag-tagged PLpro-TM from PEDV or IBV and HA-Importin α1-Myc in HEK-293T cells. As shown in Fig 3A-3B, immunoprecipitation of Flag-PLpro-TM co-precipitated exogenous expressed Importin α1 and conversely, pull-down of exogenous expressed Importin α1 (with HA and Myc tag) with HA antibody captured Flag-PLpro-TM, confirming a specific interaction between PLpro-TM and Importin α1. This interaction was conserved, as the co-immunoprecipitation were also observed between Flag-tagged PLpro-TM from MHV or PDCoV and Importin α1 (with HA and Myc tags) (S2 Fig). We further asked whether PLpro-TM interacts with endogenous Importin α1. Upon expression of Flag-tagged PLpro-TM from PEDV or IBV in HEK-293T cells, immunoprecipitation with an anti-Flag antibody consistently retrieved endogenous Importin α1 (Fig 3C). Importantly, this interaction was recapitulated during viral infection: in IBV-infected cells, an anti-nsp3 antibody co-immunoprecipitated nsp3 (which contains PLpro-TM) with endogenous Importin α1 (Fig 3D). We next examined their subcellular co-localization by immunofluorescence. In uninfected cells, Importin α1 was predominantly nuclear. Upon IBV infection, its localization shifted dynamically: at 6 h.p.i., nuclear staining diminished, while a cytoplasmic pool emerged and partially overlapped with nsp3. By 12 h.p.i., Importin α1 was distributed in both compartments, with the cytoplasmic fraction showing strong co-localization with nsp3. At 18 h.p.i., coinciding with extensive syncytium formation, Importin α1 partially re-accumulated in the nucleus, whereas the remaining cytoplasmic portion continued to co-localize with nsp3 (Fig 3E). The ER localization of PLpro-TM during infection was confirmed by co-localization of IBV nsp3 with the ER marker calnexin in infected cells (S3A Fig). Similarly, overexpressed PLpro-TM from PEDV, MHV, IBV, and PDCoV all co-localized with calnexin (S3B Fig). Collectively, these results demonstrate that IBV infection induces a dynamic redistribution of Importin α1 to the cytoplasm, likely mediated through its interaction with the ER-anchored PLpro-TM.

**Fig 3.**
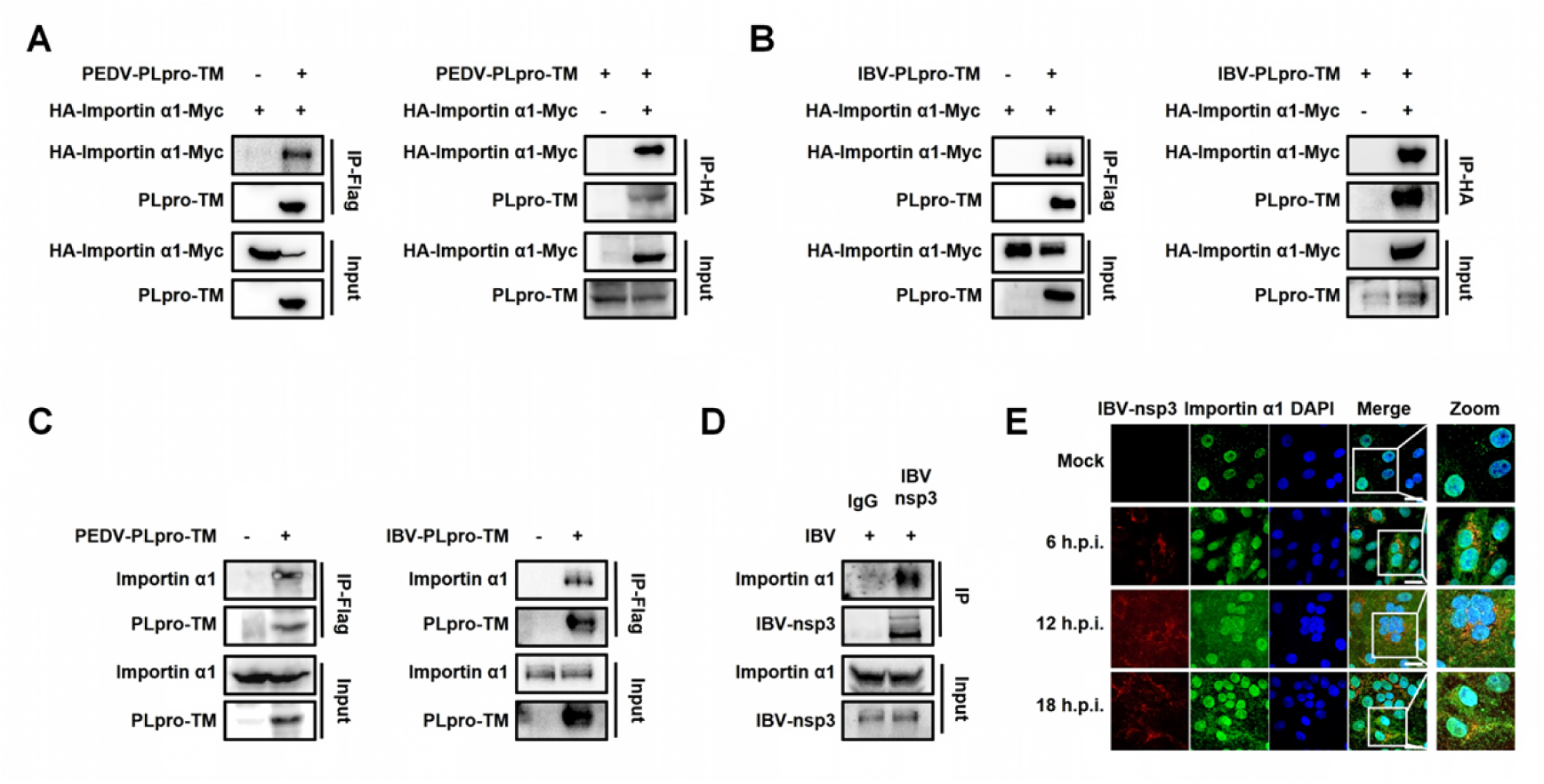
Interaction between coronavirus PLpro-TM and Importin α1. (A-B) Co-immunoprecipitation of PLpro-TM and Importin α1. Flag-tagged PLpro-TM from PEDV or IBV and Importin α1 were co-expressed in HEK-293T cells. Lysates were subjected to immunoprecipitation with an anti-Flag antibody (left panels) or an anti-HA antibody (right panels), followed by Western blot analysis. (C) Interaction of PLpro-TM with endogenous Importin α1. Flag-tagged PLpro-TM from PEDV or IBV was expressed in HEK-293T cells. Immunoprecipitation was performed with an anti-Flag antibody, and co-precipitated endogenous Importin α1 was detected by Western blot. (D) Interaction of IBV PLpro-TM with endogenous Importin α1 during infection. Lysates from mock-infected or IBV-infected cells were immunoprecipitated with an anti-nsp3 antibody, followed by Western blot analysis for the IBV nsp3 and endogenous Importin α1. (E) Co-localization of endogenous Importin α1 with IBV nsp3 during infection. Vero cells were infected with IBV or mock infected. The cells were subjected to immunostaining with antibodies against nsp3 (red) and Importin α1 (green) at 6, 12, and 18 h.p.i.. Nuclei were counterstained with DAPI (blue). Scale bar: 10 µm.

**S2 Fig.**
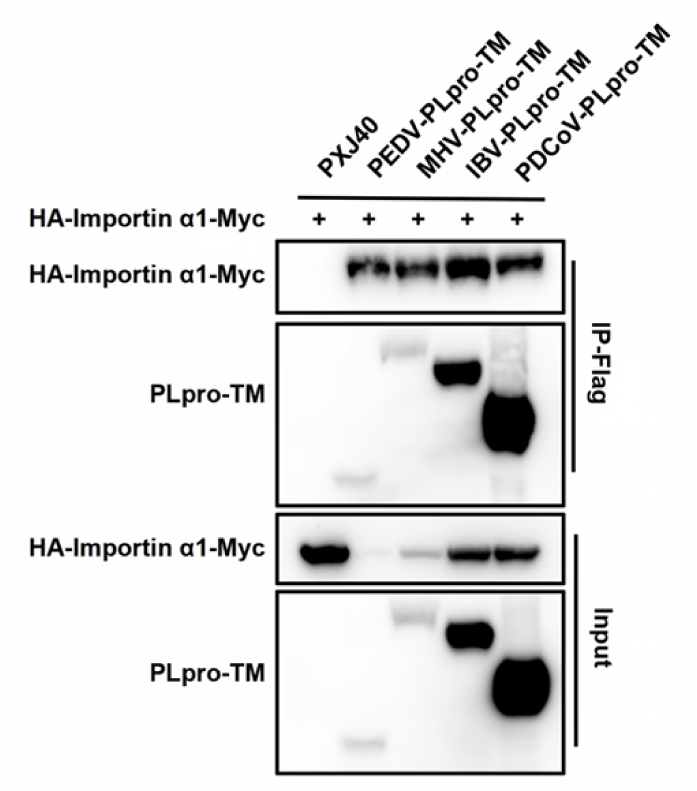
Conserved interaction between Importin α1 and PLpro-TM across coronaviruses. HA-Importin α1-Myc and Flag-tagged PLpro-TM proteins from PEDV, MHV, IBV, and PDCoV were co-expressed in HEK-293T cells. Cell lysates were subjected to immunoprecipitation using an anti-Flag antibody, followed by Western blot analysis to detect both co-precipitated HA-Importin α1-Myc and Flag-PLpro-TM.

**S3 Fig.**
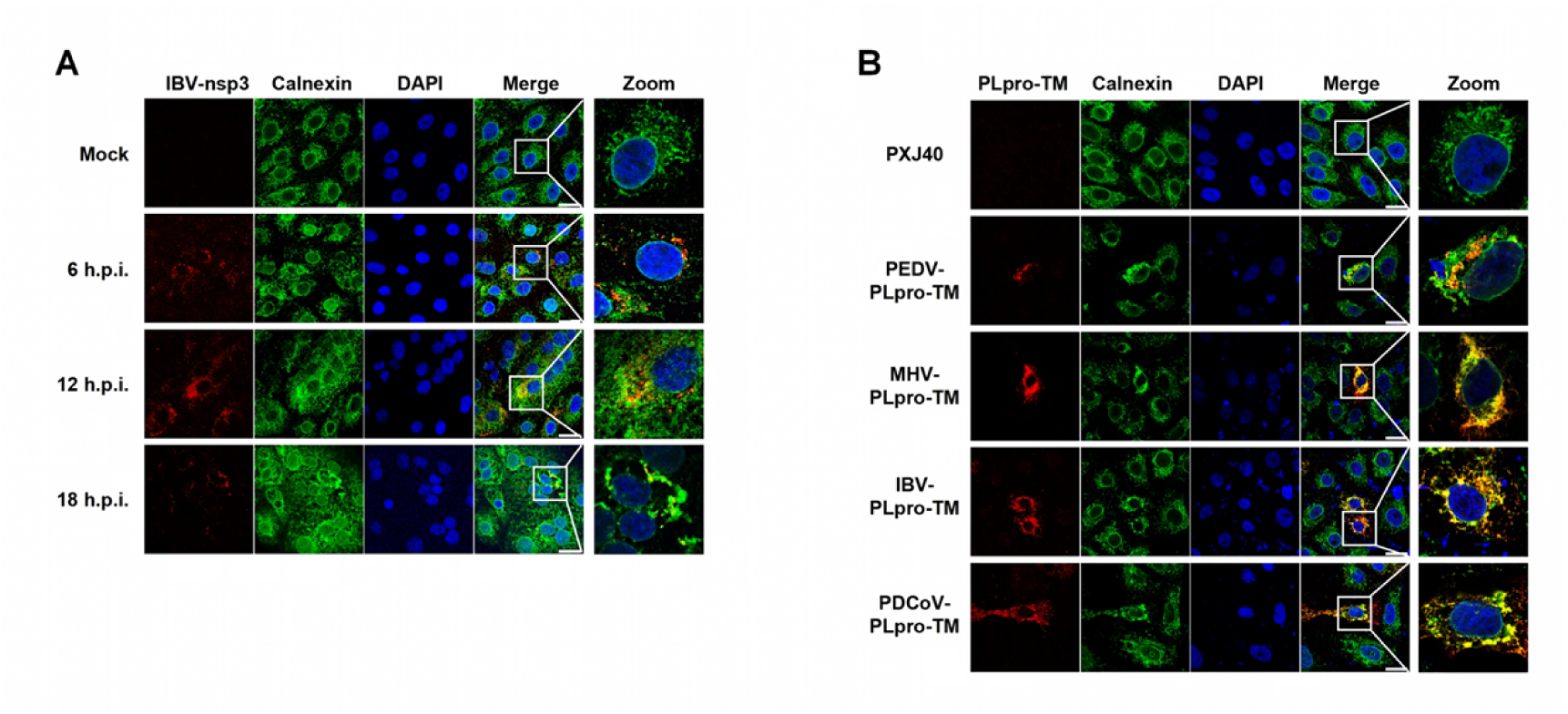
PLpro-TM localized to the endoplasmic reticulum. (A) Co-localization of IBV PLpro (located within nsp3) with ER marker calnexin during infection. Vero cells were infected with IBV or mock infected, and subjected to immunofluorescence analysis at 6, 12, and 18 h.p.i., with antibodies against IBV nsp3 (red) and the ER marker calnexin (green). Nuclei were stained with DAPI (blue). Scale bar: 10 µm. (B) Co-localization of PLpro-TM from different coronaviruses with ER marker calnexin. Vero cells were transfected with plasmids encoding Flag-tagged PLpro-TM from PEDV, MHV, IBV, or PDCoV. At 24 h.p.t., cells were fixed and co-probed with antibodies for the Flag-PLpro-TM (red) and calnexin (green). Nuclei were stained with DAPI (blue). Scale bar: 10 µm.

### Coronavirus PLpro-TM blocks Importin α1-dependent nuclear import and attenuates the expression of antiviral genes

Based on the findings that PLpro-TM interacts with Importin α1, impedes its nuclear import, and cleaves/reduces Importin α1 levels, we hypothesized that PLpro-TM could disrupt Importin α1-mediated nuclear transport. To test this, we constructed reporter plasmids containing NLS recognized by different nuclear transport receptors: SV40-NLS-GFP (with the Importin α1 and Importin α5-recognized SV40 large T antigen NLS, PKKKRKV) [8] and PY-NLS-GFP (with the Importin β2-recognized HNRNP A1-derived PY-NLS, FGYNNQSSNFGPMKGGNFGGRSSGPY) [9] (Fig 4A). Vero cells were co-transfected with either SV40-NLS-GFP or PY-NLS-GFP along with a PLpro-TM expression plasmid or vector PXJ40. As shown in the right panel of Fig 4B, both SV40-NLS-GFP and PY-NLS-GFP efficiently localized to the nucleus in control cells transfected with PXJ40. However, expression of PLpro-TM from PEDV, MHV, IBV, or PDCoV significantly inhibited the nuclear import of SV40-NLS-GFP (Importin α1-and Importin α5-dependent nuclear transport), while PY-NLS-GFP (Importin β2-dependent nuclear transport) continued to accumulate robustly in the nucleus. These results indicate that PLpro-TM specifically disrupts the Importin α-dependent nuclear import pathway without affecting transport mediated by Importin β2.

**Fig 4.**
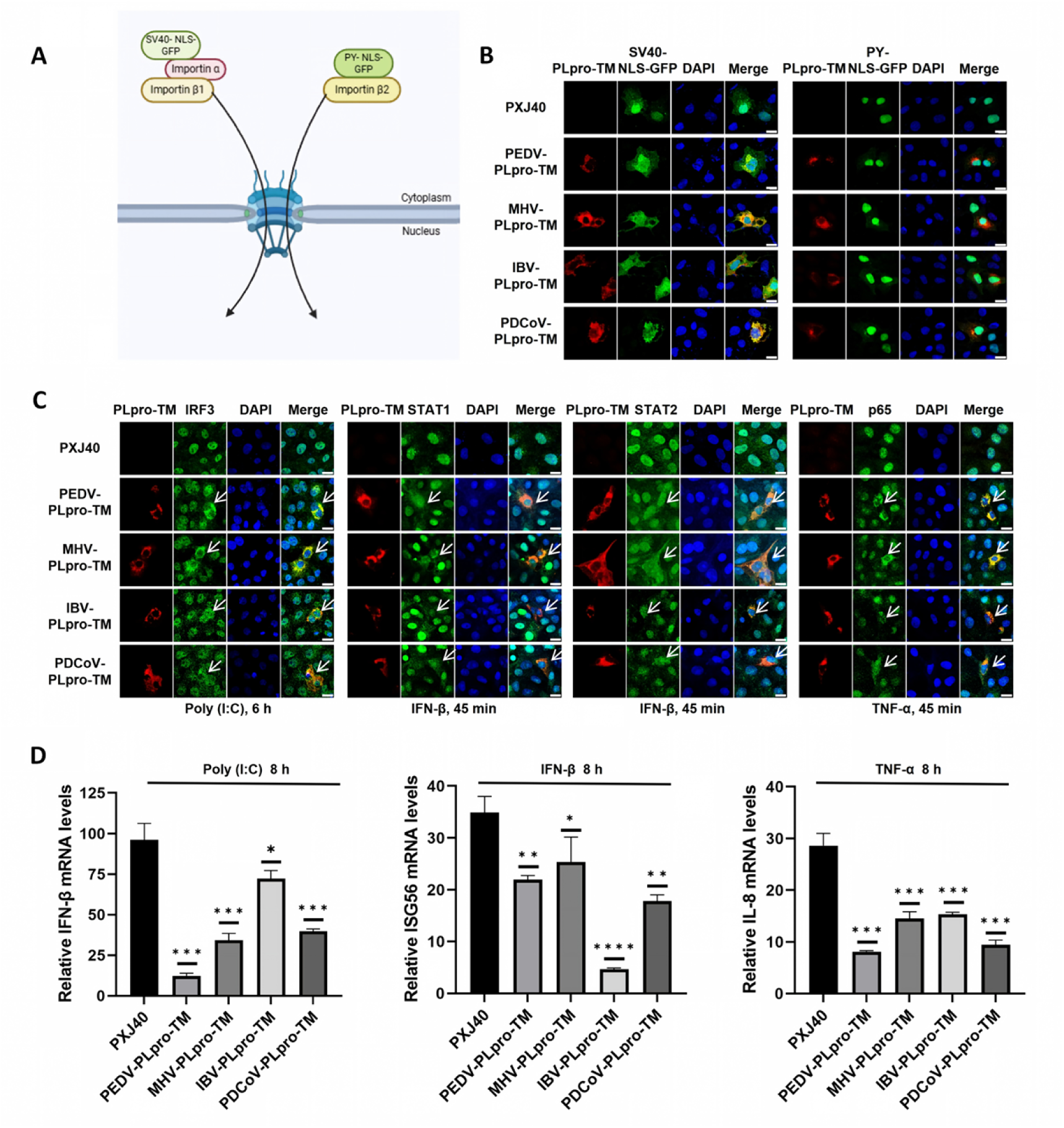
Coronavirus PLpro-TM disrupts Importin α-mediated nuclear import of transcription factors and suppresses antiviral gene expression. (A) Schematic representation of nuclear import mediated by SV40-NLS-GFP and PY-NLS-GFP. (B) Vero cells were co-transfected with plasmids encoding Flag-tagged PLpro-TM from PEDV, MHV, IBV, or PDCoV and a GFP reporter plasmid SV40-NLS-GFP and PY-NLS-GFP. Empty PXJ40 vector and GFP reporter plasmid were co-transfected as control. At 24 h.p.t, cells were processed for immunofluorescence staining to detect PLpro-TM (red) and GFP (green). Nuclei were stained with DAPI (blue). Scale bar = 10 μm. (C) Vero cells were transfected with a plasmid encoding Flag-tagged PLpro-TM from PEDV, MHV, IBV, or PDCoV or the empty PXJ40 vector At 24 h.p.t., cells were either transfected with poly(I:C) (20 μg/mL) for 6 h, treated with IFN-β (1,000 U/mL) for 45 min, or treated with TNF-α (20 ng/mL) for 45 min. Cells were then subjected to indirect immunofluorescence analysis to detect PLpro-TM (red) and endogenous IRF3, STAT1, STAT2, or p65 (green). Nuclei were stained with DAPI (blue). Scale bar = 10 μm. (D) HEK-293T cells were transfected with plasmids encoding Flag-tagged PLpro-TM from indicated coronaviruses or PXJ40 After 24 h, cells were either transfected with poly(I:C) (20 μg/mL) for 8 h, treated with IFN-β (1,000 U/mL) for 8 h, or treated with TNF-α (20 ng/mL) for 8 h. mRNA levels of IFN-β, ISG56, and IL-8 were measured by qRT-PCR. Data are presented as mean ± SD of three technical replicates from a single experiment. *P* values were calculated using Student’s t-test. *, *P* < 0.05; **, *P* < 0.01; ***, *P* < 0.001; ****, *P* < 0.0001.

Activation of the innate immune response depends critically on the Importin α-mediated nuclear translocation of key transcription factors. We therefore examined whether PLpro-TM could interfere with the nuclear import of IRF3 (NLS:^39^RTQKRLR^45^), STAT1 (NLS: ^410^KGKKTK^415^), STAT2, and p65 (NLS: ^301^RKRR^304^) - all factors that possess classical NLSs recognized by Importin α1. Vero cells were transfected with PLpro-TM from PEDV, MHV, IBV, or PDCoV, and stimulated with poly(I:C), IFN-β, or TNF-α. As shown in Fig 4C, poly(I:C), IFN-β, or TNF-α potently induced the nuclear accumulation of their respective transcription factors IRF3, STAT1, STAT2, and p65. In contrast, PLpro-TM expression resulted in the cytoplasmic retention of these transcription factors (indicated by white arrows). These data demonstrate that PLpro-TM broadly disrupts the Importin α-mediated nuclear translocation of immune transcription factors.

To determine whether this PLpro-TM–mediated blockade of nuclear import functionally impacts downstream gene expression, we measured transcription levels of key antiviral genes. qRT-PCR analysis revealed that stimulation with poly(I:C), IFN-β, or TNF-α strongly induced expression of IFN-β (dependent on nuclear entry of IRF3), ISG56 (dependent on nuclear entry of STAT1/STAT2/IRF9), or IL-8 (dependent on nuclear entry of p65), respectively; however, expression of PLpro-TM from PEDV, MHV, IBV, and PDCoV significantly reduced mRNA levels of IFN-β, ISG56, and IL-8 to varying degrees (Fig 4D). These findings indicate that PLpro-TM suppresses innate immune signaling by inhibiting the Importin α–mediated nuclear translocation of IRF3, STAT1, STAT2, and p65, thereby attenuating the expression of downstream antiviral genes.

### The protease activity and transmembrane domain of PLpro-TM are essential for downregulating Importin α1 and disrupting Importin α1-dependent nuclear transport

Coronavirus PLpro is a cysteine protease that depends on a conserved catalytic triad (Cys-His-Asp) for its enzymatic activity, which is essential for both viral polyprotein 1a and 1ab processing and deubiquitinating/de-ISGylating functions [55, 61]. To investigate the contribution of PLpro catalytic activity and its transmembrane domain to Importin α1–mediated nuclear transport, we generated catalytic-deficient mutants of PEDV-PLpro-TM (C99A, H258A, and D271A), TM-truncated PEDV-PLpro, IBV-PLpro-TM (C101A, H264A, and D275A), and TM-truncated IBV-PLpro (Fig 5A) [62, 63]. Co-expression of wild-type or mutant PLpro-TM with Importin α1 revealed differential effects on Importin α1 stability (Fig 5B-5C). Wild-type PLpro-TM strongly reduced Importin α1 levels, whereas PEDV PLpro-TM-C99A and PLpro-TM-H258A, IBV PLpro-TM-C101A and PLpro-TM-H264A, failed to do so, indicating that the enzymatic center Cys and His residues are essential for reducing Importin α1. It was noted that the PEDV PLpro-TM D271A and IBV PLpro-TM-D275A mutants degraded Importin α1 as effectively as the wild-type PLpro-TM, consistent with the report that Asp does not directly participating in hydrolysis [51, 61]. Notably, TM-truncated PEDV PLpro and IBV PLpro lost the ability to reduce Importin α1 expression, highlighting the essential role of the TM in helping PLpro reduce Importin α1 level.

**Fig 5.**
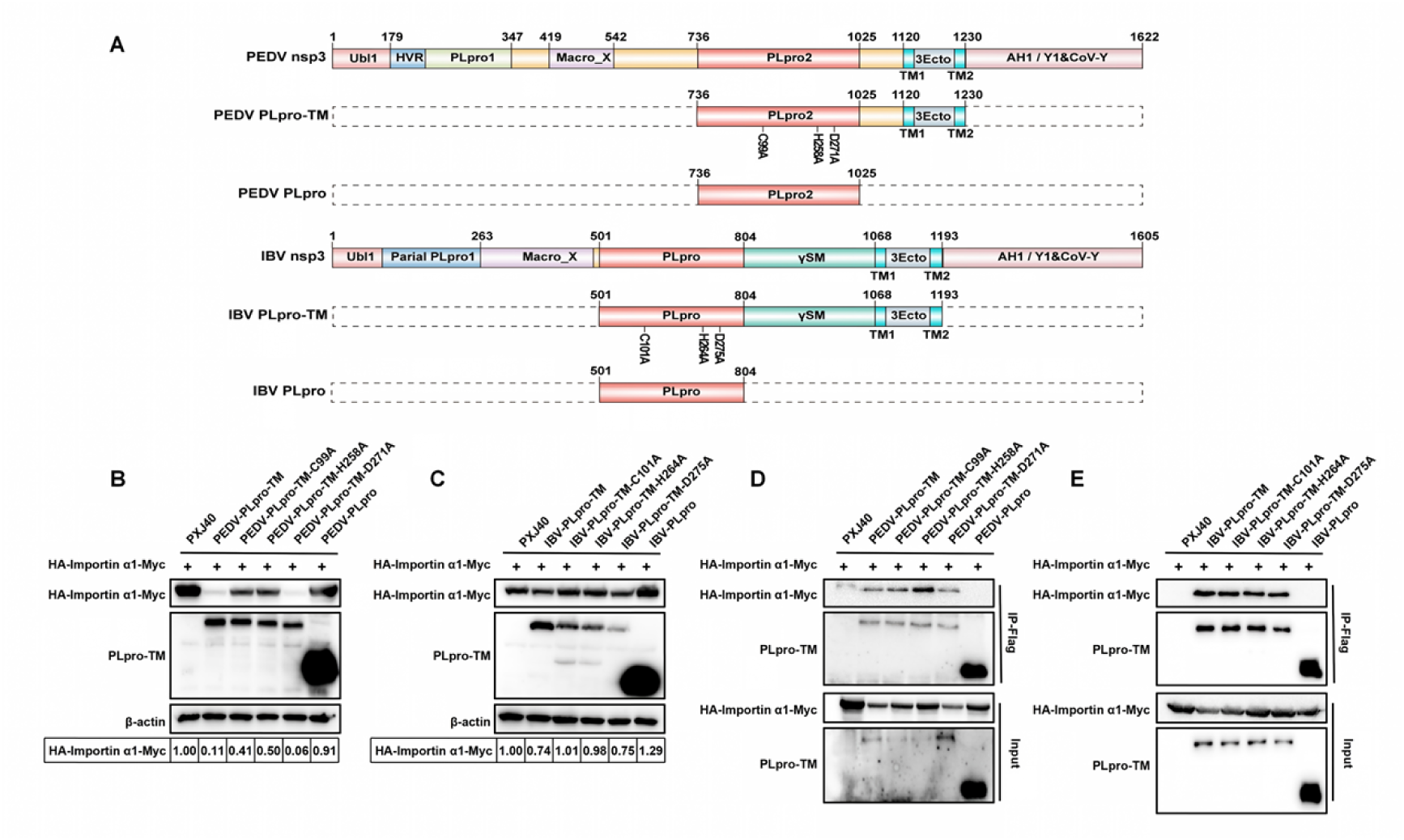
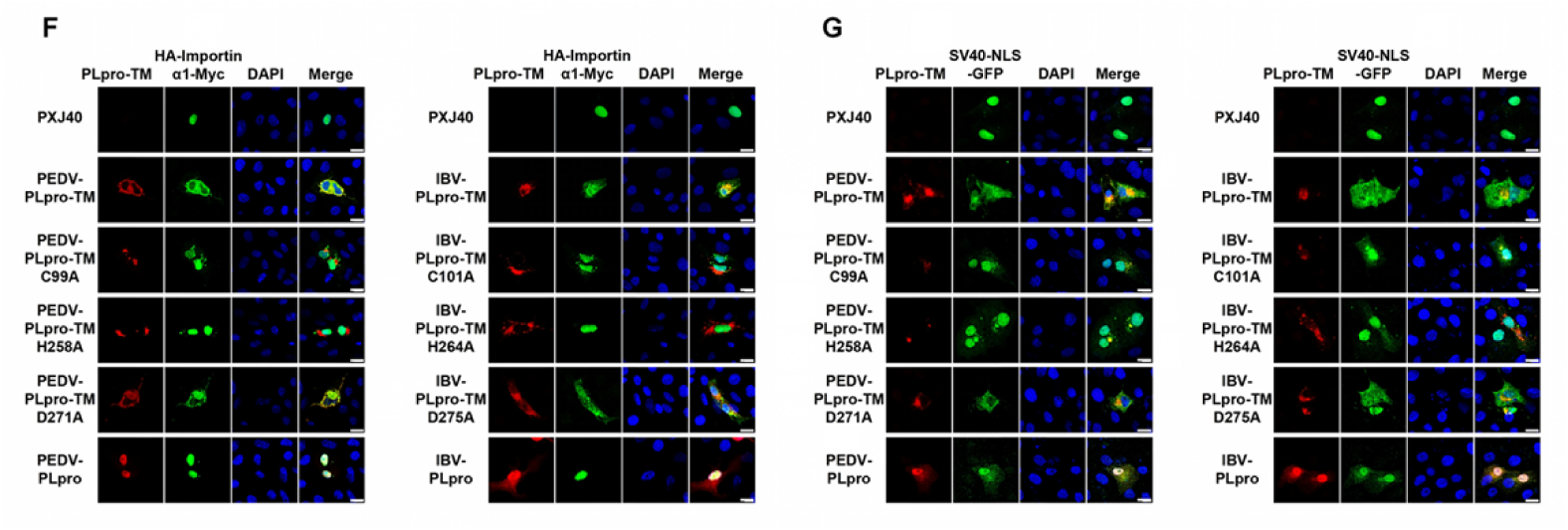
PLpro-TM enzymatic activity and TM are indispensable for downregulating Importin α1 and disrupting nuclear import function. (A) Schematic representation of PEDV/IBV PLpro-TM and their mutants. (B) Flag-tagged PEDV PLpro-TM or its mutants (C99A, H258A, D271A and PLpro) were co-expressed with Importin α1 (tagged with HA and Myc) in HEK-293T cells and subjected to Western blot analysis. (C) Flag-tagged IBV PLpro-TM or its mutants (C101A, H264A, D275A and PLpro) were co-expressed with Importin α1 (tagged with HA and Myc) in HEK-293T cells and subjected to Western blot analysis. (D) Flag-tagged PEDV PLpro-TM or its mutants and Importin α1 (tagged with HA and Myc) were co-expressed in HEK-293T cells. Cell lysates were subjected to immunoprecipitation with Anti-Flag antibody, and analyzed with Western blot. (E) Flag-tagged IBV PLpro-TM or its mutants and Importin α1 (tagged with HA and Myc) were co-expressed in HEK-293T cells. Cell lysates were subjected to immunoprecipitation with Anti-Flag antibody, and analyzed with Western blot. (F) Flag-tagged PEDV PLpro-TM or its mutants, Flag-tagged IBV PLpro-TM or its mutants, were co-expressed with Importin α1 (tagged with HA and Myc) in Vero cells. At 24 h.p.t., immunofluorescence staining was performed to detect Flag-tagged PLpro-TM (red) and HA-Importin α1-Myc (green). Cell nuclei were stained with DAPI (blue). Scale bar = 10 μm. (G) Flag-tagged PEDV PLpro-TM or its mutants, Flag-tagged IBV PLpro-TM or its mutants, were co-expressed with SV40-NLS-GFP in Vero cells. At 24 h.p.t., immunofluorescence staining was performed to detect Flag-tagged PLpro-TM (red) and GFP (green). Cell nuclei were stained with DAPI stain (blue). Scale bar = 10 μm.

**S4 Fig.**
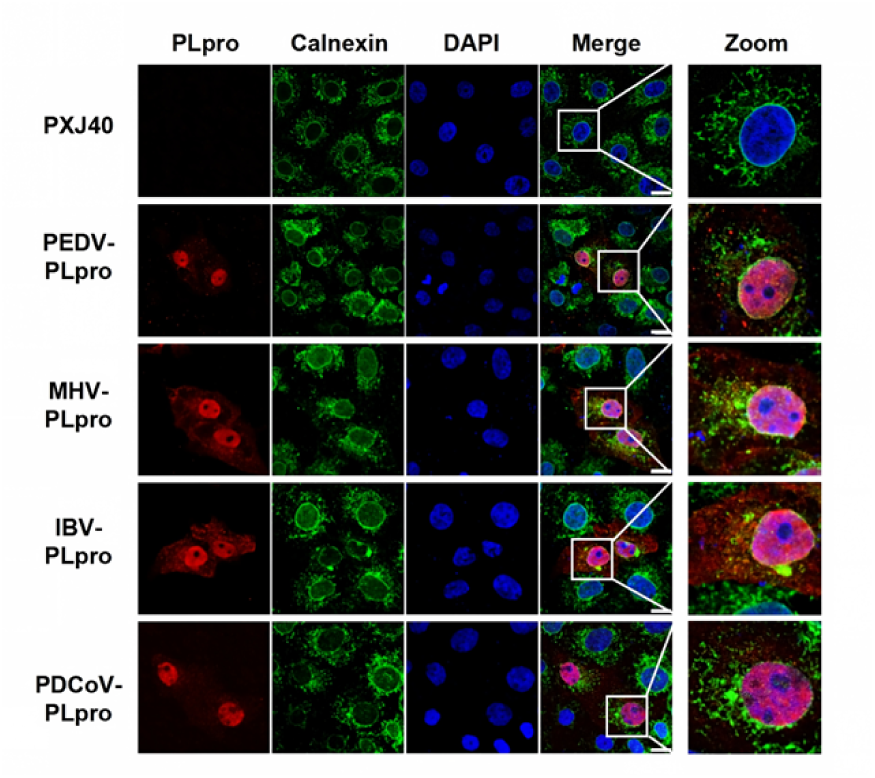
Subcellular Localization of PLpro truncated with transmembrane domain transmembrane domain. Vero cells were transfected with plasmids encoding Flag-tagged PLpro from PEDV, MHV, IBV, or PDCoV. At 24 h.p.t., cells were fixed and co-stained for the Flag-PLpro (red) and calnexin (green). Nuclei were stained with DAPI (blue). Scale bar: 10 µm.

Next, we asked whether the enzymatic activity and TM of PLpro-TM are required for its interaction with Importin α1. Flag-tagged PEDV PLpro-TM, IBV PLpro-TM, or their mutants were co-expressed with Importin α1, and co-immunoprecipitation assay was performed using anti-Flag antibody. As shown in Fig 5D-5E, both wild-type and catalytic mutants of PEDV and IBV PLpro-TM co-precipitated efficiently with Importin α1. However, both TM-truncated PEDV PLpro and IBV PLpro exhibited no binding with Importin α1. These results indicate that the TM mediated ER localization, but not the catalytic activity, is indispensable for helping the physical interaction between PLpro-TM and Importin α1.

To assess the impact of PLpro-TM enzymatic activity and its TM-mediated ER localization on the subcellular localization of Importin α1, we performed immunofluorescence assay. As shown in the left panels of Fig 5F–5G, Importin α1 exhibited predominant nuclear localization when expressed alone. In contrast, co-expression with wild-type PEDV or IBV PLpro-TM led to a marked cytoplasmic redistribution of Importin α1, which extensively colocalized with PLpro-TM. A similar re-localization of Importin α1 was observed with the catalytically impaired D271A (PEDV) and D275A (IBV) mutants, indicating that these variants retained the ability to sequester Importin α1 in the cytoplasm. However, the C99A and H258A mutants of PEDV PLpro-TM, as well as the C101A and H264A mutants of IBV PLpro-TM, showed a diminished capacity to inhibit nuclear import of Importin α1, as a substantial portion of Importin α1 was detected in the nucleus despite partial cytoplasmic colocalization. Notably, deletion of the TM disrupted the ER localization of PLpro, leading to nuclear accumulation of PLpro (S4 Fig). This loss of membrane anchoring of PLpro fully restored the nuclear localization of Importin α1 (Fig 5F), indicating that ER localization is essential for PLpro-TM to interact with and retain Importin α1 in the cytoplasm.

To functionally assess the effect of PLpro-TM enzymatic activity and its TM-mediated ER localization on Importin α1-dependent nuclear import, we employed the SV40-NLS-GFP reporter system. As shown in the right panels of Fig 5G, co-expression of the SV40-NLS-GFP reporter with wild-type PLpro-TM or the PEDV PLpro-TM-D271A and IBV PLpro-TM-D275A mutants resulted in pronounced cytoplasmic retention of GFP, indicating impairment of nuclear import. In contrast, the PEDV PLpro-TM-C99A and PLpro-TM-H258A mutants and the IBV PLpro-TM-C101A and PLpro-TM-H264A mutants had little effect on GFP localization, which remained predominantly nuclear. Similarly, deletion of the TM fully restored nuclear import of the GFP. These results demonstrate that both catalytic integrity and ER membrane anchoring are essential for PLpro-TM to disrupt Importin α1-mediated nuclear transport. Together, our results demonstrate that PLpro-TM requires not only TM for endoplasmic reticulum targeting and subsequent Importin α1 binding, but also key catalytic residues (Cys and His), to reduce Importin α1 levels. Consequently, this mechanism effectively suppresses Importin α1-mediated nuclear transport.

### The enzymatic activity and transmembrane domain of PLpro-TM are indispensable for inhibiting the nuclear translocation of transcription factors and suppressing transcription of antiviral genes

To evaluate whether the enzymatic activity and transmembrane domain of PLpro-TM inhibiting the nuclear translocation of transcription factors, Vero cells were transfected with wild-type PLpro-TM or its mutants, subsequently stimulated with poly(I:C), IFN-β, or TNF-α to induce nuclear entry of corresponding transcription factors. As shown in Fig 6A–6B, stimulation with poly(I:C), IFN-β, or TNF-α effectively induced the nuclear translocation of the corresponding transcription factors (IRF3, STAT1/STAT2, and p65). In cells expressing wild-type PEDV PLpro-TM, mutant D271A, wild-type IBV PLpro-TM, or mutant D275A, these transcription factors were retained in the cytoplasm (white arrows), indicating that both PLpro-TM and Asp mutants retain the ability to block nuclear entry. In contrast, cells expressing the catalytic mutants PEDV C99A/H258A or IBV C101A/H264A showed predominantly nuclear localization of IRF3, STAT2, and p65 (red arrows), suggesting that these mutants have lost this inhibitory function. Notably, deletion of the TM domain in either PEDV or IBV PLpro also abolished the ability to inhibit nuclear translocation of IRF3, STAT1/STAT2, and p65 (red arrows). Furthermore, a marked reduction in STAT1 signal intensity was observed in cells expressing all PLpro-TM variants or catalytic mutants (yellow arrows), indicating that PLpro-TM specifically downregulates STAT1 expression through a mechanism independent of its protease activity. Importantly, this STAT1-reducing effect was abrogated upon TM domain deletion, suggesting that ER membrane anchoring of PLpro is required for this function.

**Fig 6.**
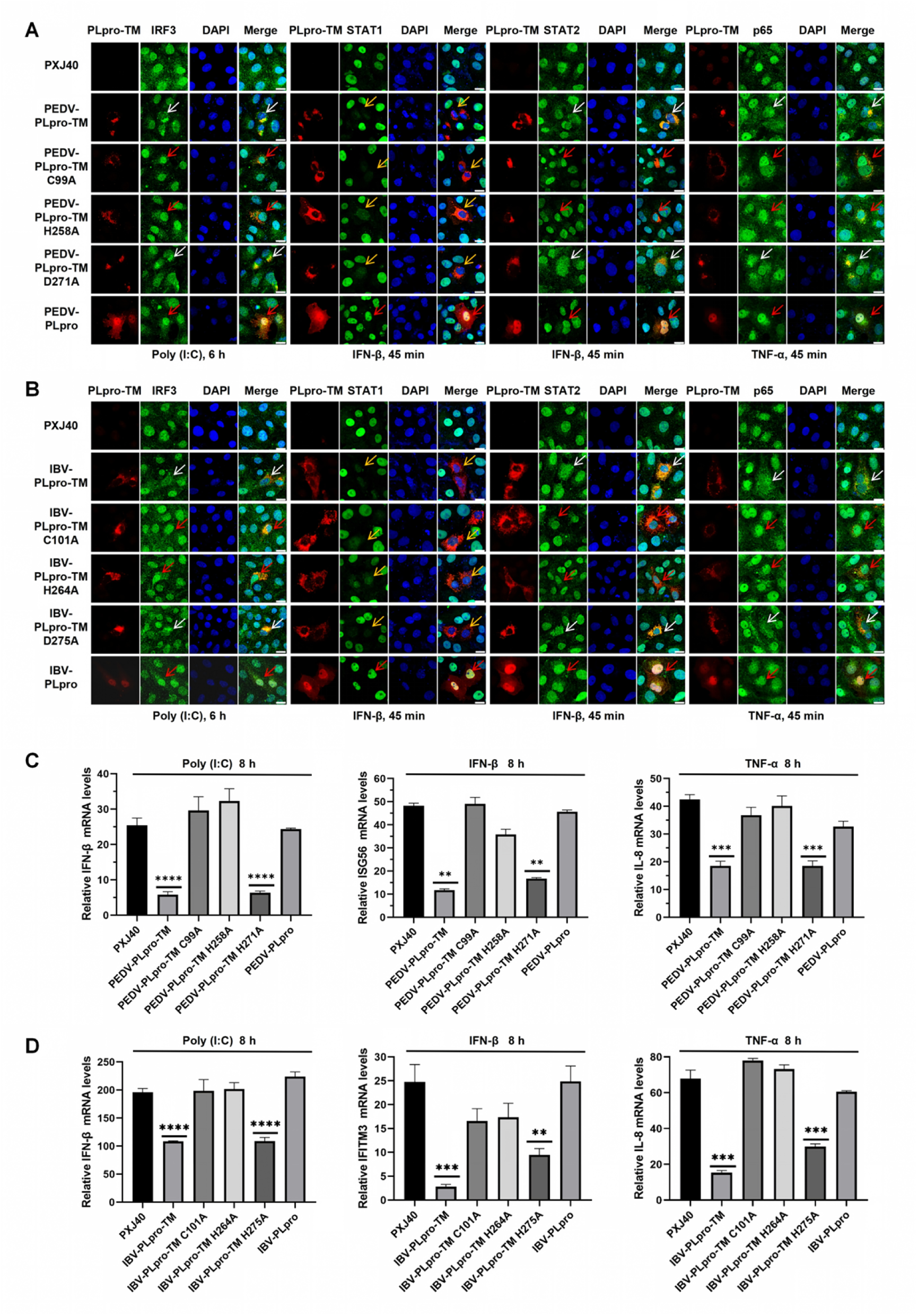
The enzymatic activity and transmembrane domain of PLpro-TM are essential for inhibiting transcription factor nuclear translocation and suppressing antiviral gene transcription. (A-B) Vero cells were transfected with plasmids encoding Flag-tagged PEDV PLpro-TM or its mutants (C99A, H258A, D271A, and PLpro) (A), or with plasmids encoding Flag-tagged IBV PLpro-TM or its mutants (C101A, H264A, D275A, and PLpro) (B). The empty vector PXJ40 was transfected as a control in parallel experiments. At 24 h.p.t., cells were either transfected with poly(I:C) (20 μg/mL) for 6 h, treated with IFN-β (1,000 U/mL) for 45 min, or treated with TNF-α (20 ng/μL) for 45 min. Indirect immunofluorescence was performed to detect PLpro-TM (red), IRF3, STAT1, STAT2, or p65 (green). Cell nuclei were stained with DAPI (blue). Scale bar = 10 μm. (C-D) PK-15 cells were transfected with plasmids encoding Flag-tagged PEDV PLpro-TM or its mutants (C); DF-1 cells were transfected with plasmids encoding Flag-tagged IBV PLpro-TM or its mutants (D); the empty vector PXJ40 served as a control. At 24 h.p.t., cells were either transfected with poly(I:C) (20 μg/mL) for 8 h, treated with IFN-β (1,000 U/mL) for 8 h, or treated with TNF-α (20 ng/μL) for 8 h. The mRNA expression levels of IFN-β, ISG56, IFITM3, and IL-8 were measured by qRT-PCR. Data are presented as mean ± SD of three technical replicates from a single experiment. *P* values were calculated using Student’s t-test. **, *P* < 0.01; ***, *P* < 0.001; ****, *P* < 0.0001.

**S5 Fig.**
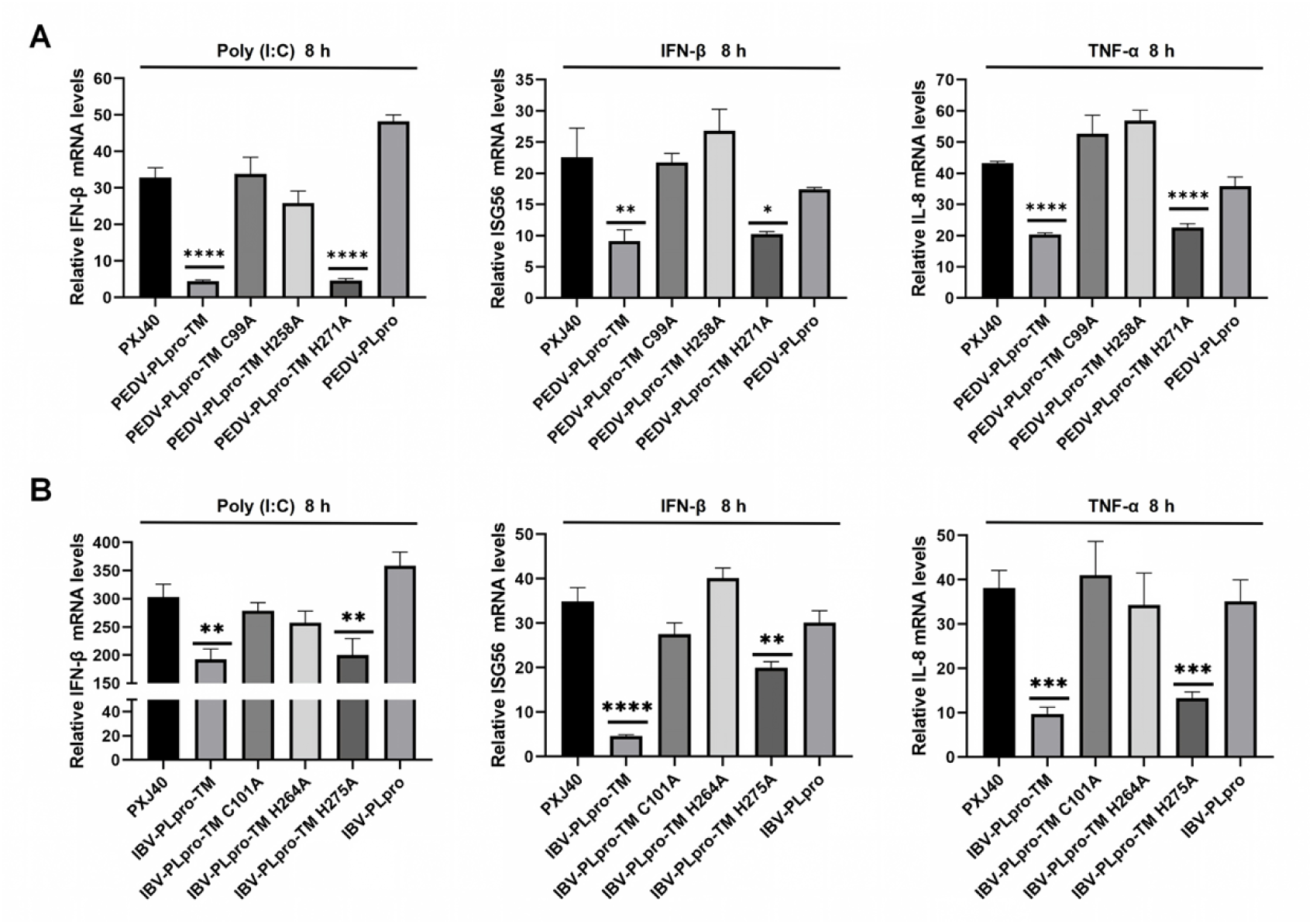
The enzymatic activity and TM domain of PLpro-TM are indispensable for suppressing transcription of antiviral genes in HEK-293T cells. (A-B) HEK-293T cells were transfected with plasmids encoding Flag-tagged PEDV PLpro-TM or its mutants (C99A, H258A, D271A, and PLpro) (A), or plasmids encoding Flag-tagged IBV PLpro-TM or its mutants (C101A, H264A, D275A, and PLpro) (B). The vector PXJ40 served as a control. At 24 h.p.t., cells were either transfected with poly(I:C) (20 μg/mL) for 8 h, treated with IFN-β (1,000 U/mL) for 8 h, or treated with TNF-α (20 ng/μL) for 8 h. The mRNA expression levels of IFN-β, ISG56, and IL-8 were measured by qRT-PCR. Data are presented as mean ± SD of three technical replicates from a single experiment. *P* values were calculated using Student’s t-test. *, *P* < 0.05; **, *P* < 0.01; ***, *P* < 0.001; ****, *P* < 0.0001.

To validate the above findings, we examined the expression of downstream genes regulated by the aforementioned transcription factors. PK-15 cells (the natural host cells of PEDV) and DF-1 cells (the natural host cells of IBV) were transfected with PEDV PLpro-TM, IBV PLpro-TM, or their respective mutants, and subsequently stimulated with poly(I:C), IFN-β, or TNF-α to induce IRF3, STAT1/STAT2, or p65 downstream gene expression. Consistent with the observed effects on transcription factor nuclear translocation in Fig 6A-6B, qRT-PCR analysis revealed that expression of PEDV PLpro-TM and the D271A mutant significantly reduced mRNA levels of IFN-β, ISG56, and IL-8 to varying degrees in PK-15 cells. In contrast, this suppressive effect was abrogated in cells expressing the catalytic mutants PEDV PLpro-TM C99A and H258A, as well as the TM-truncated PLpro mutant (Fig 6C). In DF-1 cells, expression of IBV PLpro-TM and the D275A mutant significantly reduced mRNA levels of IFN-β and IL-8 to varying degrees. Since DF-1 cells lack the ISG56 gene, IFITM3 was used as a readout for ISG expression. Both IBV PLpro-TM and the D275A mutant significantly downregulated IFITM3 transcription. In contrast, this suppressive effect was abolished in cells expressing the catalytic-null mutants C101A and H264A, as well as the TM-truncated PLpro mutant (Fig 6D). The above results were further validated in HEK-293T cells (S5 Fig). Together, these results demonstrate that both the key catalytic residues (Cys and His) and the transmembrane domain of PLpro-TM are essential for inhibiting the nuclear translocation of transcription factors and suppressing the expression of downstream antiviral genes.

### PEDV and IBV PLpro-TM cleave Importin α1 at glycine 129, whereas MHV and PDCoV PLpro-TM target Importin α1 at glycine 119

To identify the cleavage site of PLpro-TM on Importin α1, we first performed a sequence-based prediction using WebLogo (http://weblogo.threeplusone.com/), based on the polyprotein sequences known to be cleaved by PLpro of PEDV, MHV, IBV, and PDCoV [64]. This analysis revealed that coronavirus PLpro-TM exhibits a strong preference for a glycine residue at the P1 position (Fig 7A). A weak cleavage product of approximately 45 kDa was detected by an anti-Myc antibody when the 60 kDa Importin α1 (with Myc-tag at the C-terminus) was co-expressed with PLpro-TM (Fig 2D). Based on the size of this fragment, we hypothesized that the cleavage event occurs near the N-terminus of Importin α1 and predicted five potential P1 cleavage sites within this region: G72, G84, G119, G129, and G162. To evaluate these candidate sites, we constructed a series of Importin α1 mutants, each with a single glycine (G) substituted by alanine (A) at the predicted positions (Fig 7B). These mutants were then co-transfected with PLpro-TM into HEK293T cells to specifically examine their cleavage by PLpro-TM. As shown in the left panel of Fig 7C, co-expression of Importin α1 and its mutants with the empty vector PXJ40 did not alter the expression levels of wild-type Importin α1 or its mutants. In contrast, co-expression with IBV, MHV, or PDCoV PLpro-TM resulted in the appearance of a distinct 45 kDa cleavage fragment of Importin α1. Notably, when co-expressed with PEDV PLpro-TM, the full-length Importin α1 levels were markedly reduced, yet the 45 kDa cleavage product was not detected (Fig 7C), suggesting rapid degradation of the cleaved fragment. Assessment of the Importin α1 mutants revealed that, unlike mutations at other glycine residues (G72, G84, G119, G162), the G129A mutation led to a significant accumulation of full-length Importin α1 when co-expressed with PEDV PLpro-TM, and G129A mutation resulted in the disappearance of the 45 kDa cleavage product when co-expressed with IBV PLpro-TM (Fig 7C). These results suggest that the proteolytic efficiency of both PEDV and IBV PLpro-TM on Importin α1 is specifically compromised when G129 is mutated. MHV and PDCoV PLpro-TM-mediated cleavage of Importin α1 (generating the 45 kDa fragment) was unaffected by the G72A, G84A, G129A, or G162A substitutions, but was specifically abolished by the G119A mutation. Taken together, these findings indicate that PEDV and IBV PLpro-TM cleave Importin α1 at G129, whereas MHV and PDCoV PLpro-TM cleave Importin α1 at G119.

**Fig 7.**
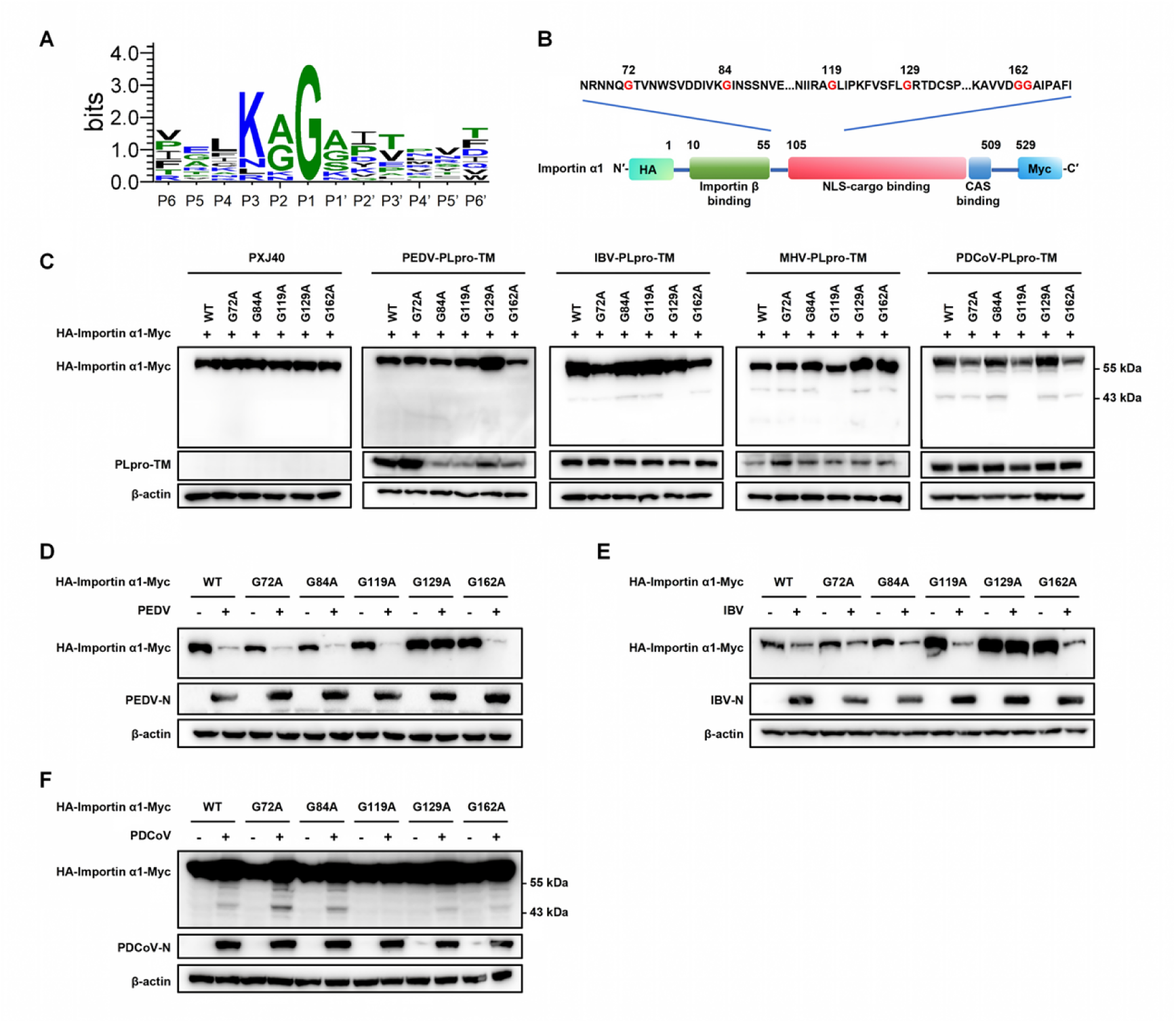
PEDV and IBV PLpro-TM cleave Importin α1 at residue G129, whereas MHV and PDCoV PLpro-TM target residue G119. (A) WebLogo analysis of the cleavage sites of PLpro based on the polyprotein 1a and 1ab cleavage sequences LXGG and other known substrates. (B) Schematic representation of Importin α1 and its mutants. (C) HEK-293T cells were co-transfected with plasmids encoding Flag-tagged PLpro-TM proteins from PEDV, MHV, IBV, or PDCoV, along with either wild-type Importin α1 or the indicated Importin α1 mutants (G72A, G84A, G119A, G129A, and G162A). At 24 h.p.t., cell lysates were prepared and analyzed by Western blotting. (D–F) Vero cells or LLC-PK1 cells were transfected with plasmids encoding wild-type Importin α1 or the indicated mutants for 24 h. At 24 h.p.t., Vero cells were infected with PEDV (D) or IBV (E) at an MOI of 5, and LLC-PK1 cells were infected with PDCoV (F) at an MOI of 5. At 18 h.p.i., Importin α1 cleavage was assessed by Western blotting with anti-Myc, and viral infection was verified using anti-N antibodies for PEDV, IBV, and PDCoV. Actin served as a loading control.

To validate our findings in the context of viral infection, we overexpressed either wild-type or mutant Importin α1 in Vero cells and then infected the cells with PEDV, IBV, or PDCoV. As shown in Fig 7D–7E, PEDV and IBV infection significantly reduced the protein levels of Importin α1 as well as the G72A, G84A, G119A, and G162A mutants. In contrast, the G129A mutant exhibited a marked accumulation of full-length Importin α1 and remained resistant to cleavage during infection. Notably, the anticipated 45 kDa cleavage fragment was not detected in PEDV- or IBV-infected cells, likely due to rapid degradation following cleavage. Similarly, PDCoV infection of LLC-PK1 cells induced cleavage of Importin α1 and the G72A, G84A, G129A, and G162A mutants, with the appearance of a 45 kDa cleavage fragment. In contrast, the G119A mutant was completely resistant to cleavage, as evidenced by the absence of the 45 kDa band (Fig 7F). These results further confirm that Importin α1 is cleaved at G129 during PEDV and IBV infection and at G119 during PDCoV infection.

### The cleavage-resistant mutations (G119A and G129A) in Importin α1 restore nuclear import function and promote anti-viral gene expression

To assess whether the cleavage-resistant G129A or G119A mutant of Importin α1 restores nuclear import ability, we co-transfected Vero cells with either PEDV or IBV PLpro-TM together with the G129A mutant or wild-type Importin α1, or with MHV or PDCoV PLpro-TM together with the G119A mutant or wild-type Importin α1. Indirect immunofluorescence analysis revealed that both wild-type Importin α1 and the cleavage-resistant mutants (G129A or G119A) were predominantly nuclear when expressed alone (Fig 8A). However, in PLpro-TM-expressing cells, in contrast to the cytoplasmic accumulation of wild-type Importin α1, the G129A and G119A mutants were able to translocate into the nucleus (Fig 8A), indicating that the cleavage-resistant mutants retain their nuclear translocation ability in the presence of PLpro-TM. These results suggest that PLpro-TM-mediated cleavage of Importin α1 is both necessary and sufficient to disrupt its nuclear import function, leading to its cytoplasmic accumulation. The above results demonstrate that the G129A and G119A mutants retain their nuclear import ability in the presence of PLpro-TM expression. We therefore further investigated whether these cleavage-resistant mutants could restore the nuclear transport of cargo proteins under PLpro-TM expression. The Importin α-dependent SV40-NLS-GFP reporter was co-transfected into Vero cells along with either wild-type Importin α1 or the G129A mutant together with PLpro-TM (PEDV or IBV), or wild-type Importin α1 or the G119A mutant together with PLpro-TM (MHV or PDCoV). As shown in Fig 8B, co-expression of the SV40-NLS-GFP reporter with wild-type Importin α1 and PLpro-TM resulted in pronounced cytoplasmic retention of GFP, indicating an impairment of nuclear import. In contrast, co-expression of the GFP reporter with the Importin α1 G129A or G119A mutant and PLpro-TM led to nuclear localization of GFP, suggesting that the cleavage-resistant mutant G129A and G119A mutants restore nuclear import ability in the presence of PLpro-TM.

**Fig 8.**
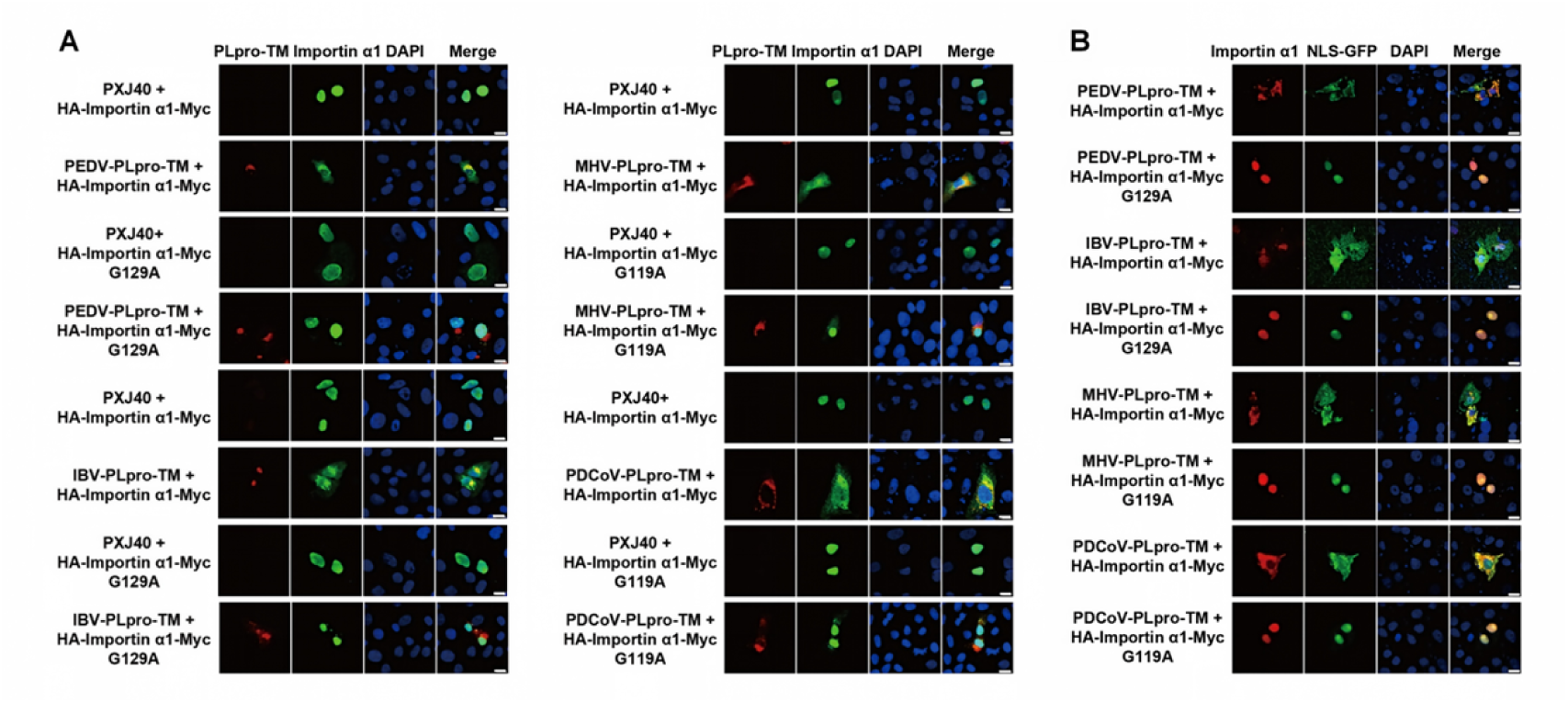
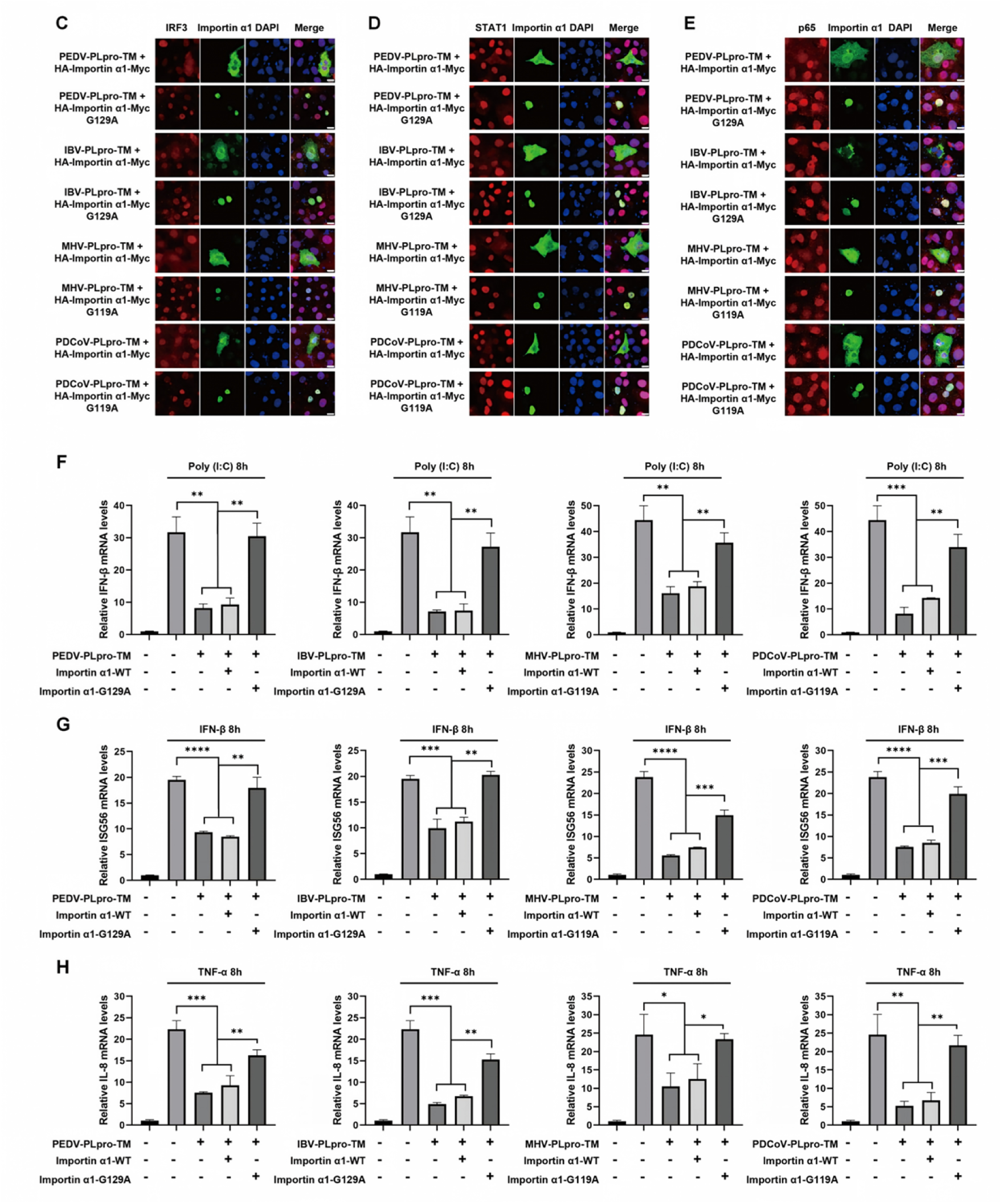
Cleavage-resistant mutation (G119A or G129A) in Importin α1 restores nuclear import ability and IFN-β signaling, as well as NF-κB signaling. (A) Vero cells were co-transfected with plasmids encoding wild-type Importin α1 or its G129A or G119A mutant (with HA tag at N terminus and Myc tag at C terminus), along with Flag-tagged PLpro-TM from PEDV, MHV, IBV, or PDCoV, or the empty vector PXJ40. At 24 h.p.t., immunofluorescence staining was performed to detect Flag-tagged PLpro-TM (red) and Importin α1 or its mutants (green). Cell nuclei were labeled with DAPI (blue). Scale bar = 10 μm. (B) Vero cells were co-transfected with the SV40-NLS-GFP reporter along with plasmids encoding wild-type Importin α1 or its G129A or G119A mutant, together with Flag-tagged PLpro-TM from PEDV, MHV, IBV, or PDCoV. At 24 h.p.t., immunofluorescence was performed to detect Importin α1 or its mutants (red) and GFP (green). Cell nuclei were labeled with DAPI (blue). Scale bar = 10 μm. (C-E) Vero cells were co-transfected with plasmids encoding wild-type HA-Importin α1-Myc or its G129A or G119A mutant, along with Flag-tagged PLpro-TM from PEDV, MHV, IBV, or PDCoV. At 24 h.p.t., cells were either transfected with poly(I:C) (20 μg/mL) for 6 h (C), or treated with IFN-β (1,000 U/mL) for 45 min (D), or treated with TNF-α (20 ng/μL) for 45 min (E). Indirect immunofluorescence was performed to detect Importin α1 or its mutants (green) and IRF3 (red) (C), STAT1 (red) (D), or p65 (red) (E). Cell nuclei were stained with DAPI (blue). Scale bar = 10 μm. (F-H) HEK-293T cells were co-transfected with plasmids encoding wild-type HA-Importin α1-Myc, or its G129A or G119A mutant, or pCMV-Myc vector, together with Flag-tagged PLpro-TM from PEDV, MHV, IBV, or PDCoV. At 24 h.p.t., cells were either transfected with poly(I:C) (20 μg/mL) for 8 h (F), or treated with IFN-β (1,000 U/mL) for 8 h (G), or treated with TNF-α (20 ng/μL) for 8 h (H). The mRNA expression levels of IFN-β (F), ISG56 (G) and IL-8 (H) were measured by qRT-PCR. Data are presented as mean ± SD of three technical replicates from a single experiment. *P* values were calculated using Student’s t-test. **, *P* < 0.01; ***, *P* < 0.001; ****, *P* < 0.0001.

We further assessed whether these cleavage-resistant Importin α1 mutants could restore the nuclear transport of transcription factors and the expression of antiviral genes in PLpro-TM-expressing cells. Vero cells were co-transfected with either wild-type Importin α1 or its G129A mutant together with PLpro-TM (PEDV or IBV), or with wild-type Importin α1 or its G119A mutant together with PLpro-TM (MHV or PDCoV). Subsequently, the cells were stimulated with poly(I:C), IFN-β or TNF-α to induce the nuclear translocation of IRF3, STAT1 and p65, as well as the expression of IFN-β, ISG56 and IL-8. Immunofluorescence analysis revealed that in cells expressing wild-type Importin α1 and PLpro-TM (Plasmids encoding HA-Importin α1-Myc and Flag-tagged PLpro-TM were co-transfected into the same cells), IRF3, STAT1 and p65 accumulated in the cytoplasm upon corresponding stimulation (Fig 8C–8E), and the expression of IFN-β, ISG56 and IL-8 was significantly suppressed, as assessed by qRT-PCR (Fig 8F–8H). In contrast, in cells expressing the G129A or G119A mutant together with PLpro-TM (Plasmids encoding HA-Importin α1-Myc G129A or G119A mutants and Flag-tagged PLpro-TM were co-transfected into the same cells), IRF3, STAT1 or p65 was efficiently translocated into the nucleus alongside the G129A or G119A mutant (Fig 8C–8E), and the expression of IFN-β, ISG56 and IL-8 was restored (Fig 8F–8H). These results demonstrate that, in cells expressing PLpro-TM, mutating glycine 129 or glycine 119 of Importin α1 to alanine confers resistance to cleavage, thereby functionally rescuing nuclear import capability and restoring IFN-β signaling and NF-κB signaling. This suggests that functional Importin α1 (cleavage-resistant) helps the host cell initiate antiviral responses.

We next evaluated the functional impact of Importin α1 cleavage by PLpro-TM on coronavirus replication. As shown in Fig 9A–9C, overexpression of wild-type HA-Importin α1-Myc slightly inhibited PEDV, IBV, and PDCoV replication, as evidenced by reduced viral N protein levels in cells (determined by Western blotting) and reduced virus release in the supernatant (determined by qRT-PCR targeting the N gene). Notably, the cleavage-resistant mutants (G129A for PEDV/IBV and G119A for PDCoV) consistently exhibited a more potent inhibitory effect on the replication of all three coronaviruses. These results suggest that functional (cleavage-resistant) Importin α1 plays an antiviral role. Collectively, our findings demonstrate that coronavirus PLpro-TM antagonize host innate immune signaling by targeting Importin α1 for cleavage. This cleavage disrupts Importin α1-mediated nuclear transport, thereby abolishing its antiviral activity, suppressing downstream immune gene expression, and facilitating coronavirus infection.

**Fig 9.**
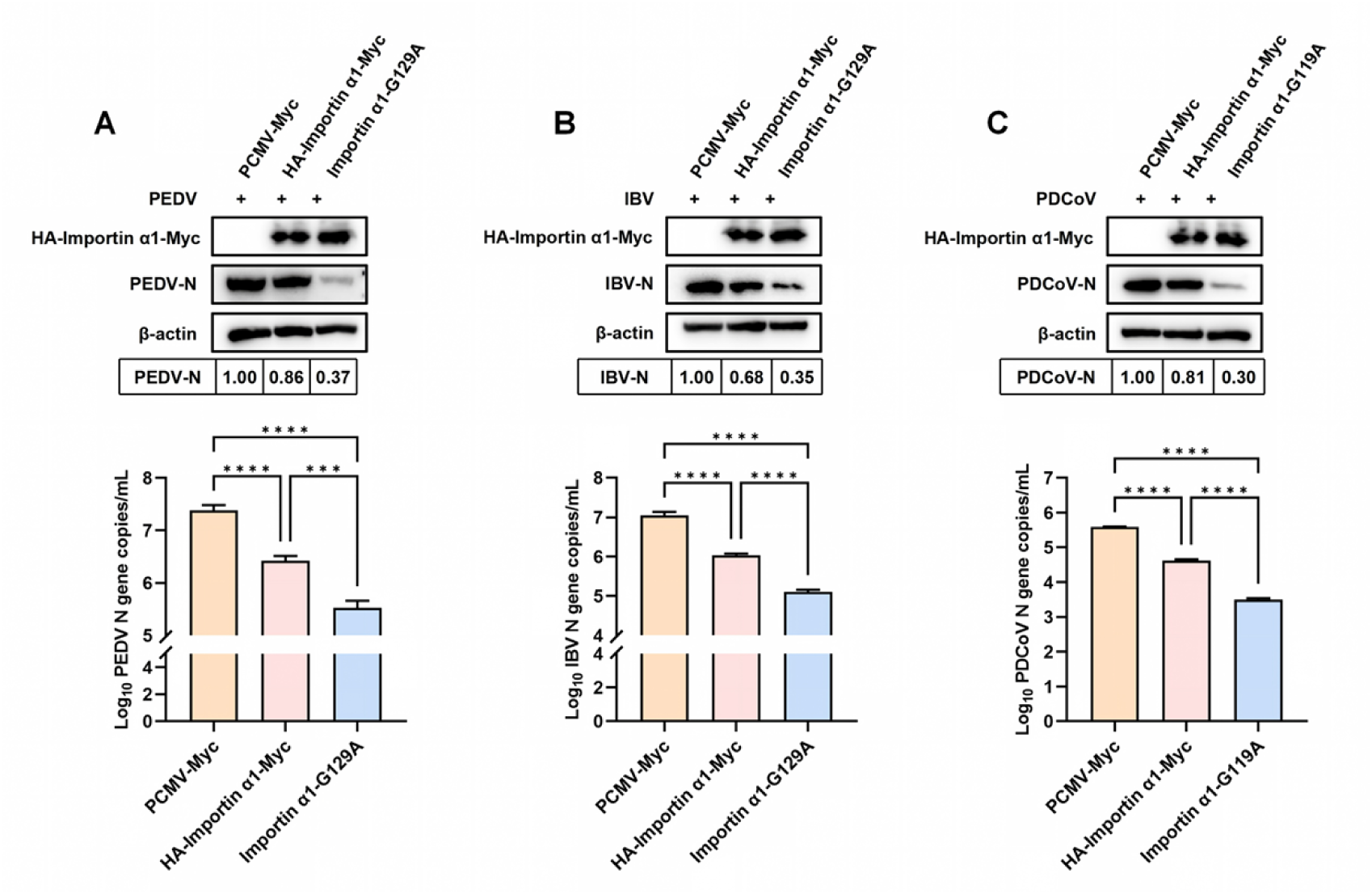
Functional (cleavage-resistant) Importin α1 suppresses PEDV, IBV and PDCoV replication. (A–C) Vero cells (A, B) or LLC-PK1 cells (C) were transfected with plasmids encoding wild-type Importin α1, the indicated cleavage-resistant mutant (G129A for panels A–B; G119A for panel C), or the empty pCMV-Myc vector for 24 h, followed by infection with PEDV (A), IBV (B), or PDCoV (C) at an MOI of 1 for 18 h. Cell lysates were analyzed by Western blotting using an anti-N antibody to confirm viral infection (upper panels). Virus particle release was assessed by quantifying N gene copy numbers in culture supernatants via qRT-PCR (lower panels). Data are presented as mean ± SD of three technical replicates from a single representative experiment. *P* values were calculated using Student’s *t*-test. ***, *P* < 0.001; ****, *P* < 0.0001.

## Discussion

In this study, we identify a previously unrecognized mechanism by which coronaviruses from all four genera (α, β, γ, δ) subvert host innate immunity through the proteolytic inactivation of Importin α1, a key receptor of nuclear import for transcription factors. We demonstrate that coronavirus encoded ER membrane-associated PLpro-TM directly cleaves Importin α1 at specific glycine residues—G129 for PEDV and IBV, and G119 for MHV and PDCoV—thereby disrupting its nuclear import function. This cleavage impairs the nuclear translocation of multiple transcription factors (IRF3, STAT1, STAT2, and p65) and suppresses the expression of downstream antiviral genes, including IFN-β, ISG56, IFITM3, and IL-8 (Fig 10). Importantly, cleavage-resistant mutants of Importin α1 (G129A or G119A) restore nuclear import capability, rescue IFN-β signaling and NF-κB signaling, and exert a more potent inhibitory effect on viral replication than wild-type Importin α1. Our work establishes PLpro-TM-mediated cleavage of Importin α1 as a conserved immune evasion strategy across coronaviruses and highlights this interaction as a potential target for broad-spectrum antiviral intervention.

**Fig 10.**
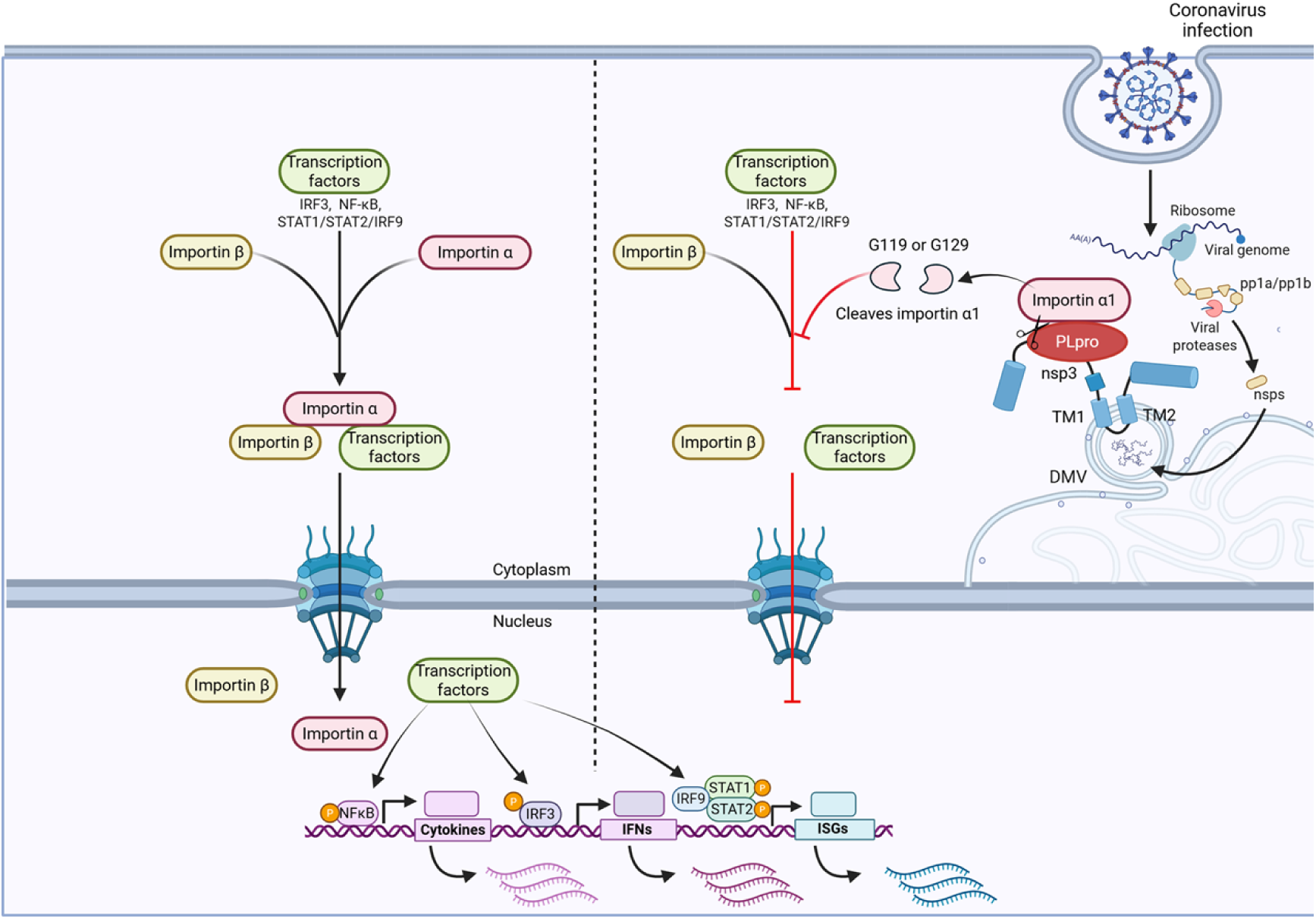
Proposed model for coronavirus PLpro-TM-induced suppression of nuclear import and antiviral immunity. The left panel depicts the classical nuclear import pathway. Cargo proteins bearing an NLS, such as transcription factors, are recognized by Importin α, which subsequently forms a ternary complex with Importin β. This Importin α/β–cargo complex is translocated through the nuclear pore complex via interactions with phenylalanine–glycine (FG)-rich nucleoporins, ultimately initiating the transcription of downstream genes. The right panel illustrates the disruption of Importin α1-mediated nuclear import by coronavirus PLpro-TM. PLpro-TM is a domain of nsp3 anchored to the ER membrane or double-membrane vesicles (DMV) via its C-terminal TM1 and TM2. The cytoplasmic-facing PLpro interacts with Importin α1 and cleaves it at residues G119 or G129, removing the N-terminal Importin β-binding (IBB) domain. This cleavage precludes the formation of the Importin α/β heterodimer, thereby abrogating the nuclear translocation of Importin α1-dependent cargo proteins, including IRF3, STAT1/2, and p65. Consequently, the expression of downstream antiviral genes is suppressed. This figure was created with BioRender (Toronto, ON, Canada).

The nuclear translocation of key signaling transcription factors is essential for initiating innate immune responses, with Importins serving indispensable roles in mediating their nuclear transport and regulatory functions [65]. It has been reported that several viruses have evolved distinct strategies to target the nuclear transport system and to subvert host defense mechanism. For instance, the NS5 protein of Japanese encephalitis virus (JEV), a member of the *Flaviviridae* family, engages multiple Importin α isoforms—including Importin α1, Importin α3, and Importin α4—thereby competitively disrupting their interaction with the transcription factors IRF3 and NF-κB p65 subunit [17]. Dengue virus NS5 contains a cryptic NLS that saturates the Importin α/β nuclear import capacity [66]. In another strategy, the VP24 protein of Ebola virus, belonging to the *Filoviridae* family, recognizes a distinct NLS-binding site on Importin α5, thus selectively blocking the nuclear import of phosphorylated STAT1 [67]. Similarly, the polymerase of hepatitis B virus, from the *Hepadnaviridae* family, interferes with STAT1/2 nuclear translocation by competitively occupying the region on Importin α5 that is essential for STAT1/2 recruitment [68]. Collectively, these examples underscore the evolutionary convergence on Importin-targeted disruption as an effective viral immune evasion strategy. Coronaviruses have also evolved several strategies to impede the nuclear translocation of transcription factors [44–48, 50]. Previous studies have focused on individual viral proteins from specific coronaviruses, each employing a distinct mechanism [20, 47–49]. Although these reports underscore the importance of targeting the nuclear transport machinery, the mechanisms described are largely viral- or protein-specific. Of note, they predominantly involve competitive inhibition of Importin–transcription factor interactions, whereas virus-mediated proteolytic cleavage of Importin has received comparatively limited attention. Our study reveals that PLpro-TM from diverse coronaviruses directly cleaves Importin α1 at glycine residues and impairs the nuclear import of transcription factors, revealing an evolutionarily conserved mechanism to escape host anti-viral response among coronaviruses. To our knowledge, this is the first report demonstrating that a viral protease directly cleaves an Importin α family member to block nuclear transport, rather than simply binding, inducing ubiquitin-proteasome-dependent degradation, or competing for NLS binding.

Coronavirus PLpro plays crucial roles in viral replication and immune evasion. As a component of nsp3, PLpro is anchored to the ER membrane via its transmembrane domain [64], where it contributes to the cleavage of newly synthesized viral polyproteins 1a and 1ab to release nsp1-nsp3, thereby providing essential elements (nsp3) for the assembly of the double membrane vesicles (DMV) for virus replication [69–72]. In addition, PLpro possesses deubiquitinating (DUB) and deISGylating activities, removing ubiquitin or ISG15 modifications from host proteins to regulate protein stability and antiviral signaling [73–75]. PLpro also directly cleaves host substrates to dysregulate cellular functions [76, 77]. A notable example is SARS-CoV-2 PLpro, which cleaves IRF3 at a canonical LGG↓G motif to suppress interferon production [76]. Here, we extend this paradigm by identifying Importin α1 as a novel host target of the PLpro-TM domain across multiple coronaviruses.

Our domain-function analysis revealed that both the catalytic activity and the transmembrane domain of PLpro-TM are essential for its ability to downregulate Importin α1 and disrupt nuclear import. Structurally, coronavirus PLpro harbors a conserved catalytic triad composed of Cys-His-Asp/Asn residues [78, 79]. We demonstrated that mutations in the catalytic residues Cys and His of PEDV and IBV PLpro-TM completely abolished its ability to cleave Importin α1, inhibit transcription factor nuclear translocation, and suppress antiviral gene expression. In contrast, mutation of the Asp residue exerted a relatively minor effect: the Asp mutant retained the capacity to block transcription factor nuclear entry and downregulate antiviral gene expression. This discrepancy likely reflects the essential catalytic roles of Cys and His in substrates hydrolysis, as mutations at these sites severely compromise or abolish enzymatic activity [80]. By comparison, the Asp residue primarily contributes to stabilizing the catalytic triad or facilitating substrate recognition, so its mutation does not completely eliminate enzymatic function [51, 61]. These results further confirm that PLpro-TM cleavage of Importin α1 relies on its classical enzymatic activity. Our findings further demonstrate that the transmembrane domain is indispensable for anchoring PLpro-TM to the ER membrane and is also critical for both its interaction with Importin α1 and the subsequent cleavage event. Mechanistically, we propose that this cleavage may occur either co-translationally during Importin α1 synthesis, or post-translationally upon its transient association with the ER membrane—a compartment spatially proximal to the ER- or DMV-anchored PLpro-TM of nsp3. Such spatiotemporal compartmentalization likely enhances the efficiency of PLpro-TM–Importin α1 engagement and facilitates the cleavage process.

Importin α1 functions as a key adaptor in nuclear transport by directly recognizing NLS-bearing cargo proteins and associating with Importin β1 to form a ternary complex, which then interacts with nuclear pore complex components to facilitate nuclear import [10, 81, 82]. Structurally, Importin α1 contains three conserved domains: an N-terminal IBB domain, a central armadillo repeat region that binds classical NLS-containing cargoes, and a C-terminal domain that interacts with the nuclear export factor CAS (also known as CSE1L) [11, 83]. Using sequence prediction and site-directed mutagenesis, we mapped the cleavage sites of Importin α1 to G129 for PEDV and IBV PLpro-TM, and to G119 for MHV and PDCoV PLpro-TM, both of which reside within the N-terminal region. Cleavage at either residue separates the IBB domain from the NLS-binding domain, thereby abolishing the ability of Importin α1 to mediate nuclear import of its cargo proteins. Although the core PLpro motif is often conserved as LXGG [64], we observed lineage-specific preferences: PEDV and IBV PLpro-TM recognize KXGG [62] and KAGG [63, 84], respectively, whereas MHV and PDCoV PLpro-TM favor LXGG [85, 86] and XXAG[61] motifs. These subtle sequence divergences might explain the differential cleavage site selection on Importin α1 (SFLG^129^↓R for PEDV/IBV PLpro-TM and IRAG^119^↓I for MHV/PDCoV PLpro-TM). The use of two distinct yet adjacent cleavage sites by different coronaviruses suggests a degree of evolutionary divergence while preserving the overall strategy of Importin α1 inactivation. Cleavage-resistant mutants (G129A and G119A) of Importin α1 not only resist proteolysis but also retain their ability to mediate nuclear import of NLS-containing cargoes, including transcription factors and a heterologous GFP reporter. These mutants restore IFN-β signaling and suppress viral replication to a significantly greater extent than wild-type Importin α1. Collectively, these gain-of-function data provide strong evidence that Importin α1 is a bona fide antiviral factor and that its cleavage by PLpro-TM represents a key pathogenic event.

In humans, multiple Importin α subtypes exist and are grouped into three subfamilies based on sequence similarity: the α1 subfamily (Importin α5/KPNA1, Importin α6/KPNA5, Importin α7/KPNA6), the α2 subfamily (Importin α1/KPNA2 and Importin α8/KPNA7), and the α3 subfamily (Importin α3/KPNA4 and Importin α4/KPNA3) [12, 13], with distinct NLS-binding affinities that confer cargo selectivity and transport specificity [87]. Importin α1 is known to mediate the nuclear translocation of transcription factors essential for antiviral immunity, including IRF3, STAT1, STAT2, and p65 [15, 20, 26]. PLpro-TM exhibits a highly selective regulatory strategy by specifically targeting Importin α1 while sparing other Importin α isoforms. This selective cleavage globally impairs the nuclear entry of key transcription factors, including IRF3, STAT1, STAT2, and p65, thereby reducing the expression of downstream effectors such as IFN-β, ISG56, IFITM3, and IL-8. Such precise targeting of Importin α1 may enable the virus to suppress critical innate immune responses while avoiding the excessive cytotoxicity or nonspecific interference that would result from broader host targeting, thus maintaining a favorable cellular environment for viral replication. In the other hand, the antagonism of Importin α1-dependent nuclear entry by PLpro-TM represents a more effective strategy than targeting individual transcription factors, and makes Importin α1 an attractive viral target. The conservation of this mechanism across multiple coronaviruses highlights its fundamental importance in viral evasion of host antiviral responses.

Several questions remain open. First, although we have identified G129 and G119 as the primary cleavage sites, the precise structural basis for substrate recognition by PLpro-TM warrants further investigation. Second, while we focused on Importin α1, our data show that PEDV PLpro-TM also moderately reduces Importin α4 levels, suggesting that additional Importin α isoforms may be targeted under certain conditions. Third, it will be important to test whether human coronaviruses, including SARS-CoV-2, SARS-CoV, MERS-CoV, HCoV-OC43, and HCoV-229E employ a similar mechanism.

In summary, our work reveals that coronavirus PLpro-TM directly cleaves Importin α1 at conserved glycine residues, thereby disrupting Importin α1-mediated nuclear transport, blocking the nuclear translocation of key antiviral transcription factors, and suppressing innate immune gene expression. This mechanism is conserved across α, β, γ, and δ coronaviruses, establishing PLpro-TM as a pan-coronavirus virulence factor and Importin α1 as a critical host antiviral node. The identification of cleavage-resistant mutants that restore antiviral function not only validates the mechanistic model but also opens the door for future therapeutic strategies aimed at preserving Importin α1 integrity during coronavirus infection. The development of small-molecule inhibitors that block PLpro-TM-mediated cleavage of Importin α1 could represent a novel broad-spectrum antiviral approach.

## Materials and Methods

### Cells and Viruses

African green monkey kidney epithelial Vero cells (CRL-1586); human embryonic kidney HEK-293T cells (CRL-3216); chicken embryo fibroblast DF-1 cells (CRL-3586), and porcine kidney-15 (PK-15) cells (CCL-33) were obtained from ATCC. Porcine kidney-1 (LLC-PK1) cells (CL-101) was kindly provided by provided by Prof. Fei Gao (Shanghai Veterinary Research Institute, CAAS, China). Vero cells, HEK-293T cells, DF-1 cells, PK-15 cells, and LLC-PK1 cells were maintained in Dulbecco’s Modified Eagle Medium (DMEM) (Gibco-Thermo Fisher, Waltham, MA, USA) supplemented with 10% (v/v) fetal bovine serum (FBS) (Gibco-Thermo Fisher, Waltham, MA, USA).

The IBV Beaudette strain was kindly provided by Prof. Dingxiang Liu’s laboratory (South China Agricultural University, China). The PEDV HLJBY strain was kindly provided by Assoc. Prof. Changchao Huan’s laboratory (Yangzhou University, China). The PDCoV strain was kindly provided by Prof. Tongling Shan’s laboratory (Shanghai Veterinary Research Institute, CAAS, China).

### Antibodies and chemicals

Mouse anti-PEDV-N monoclonal antibody was kindly provided by Prof. Yanjun Zhou (Shanghai Veterinary Research Institute, CAAS, China). Rabbit anti-IBV-N polyclonal antibody, mouse anti-IBV-N polyclonal antibody, and mouse anti-IBV-nsp3 polyclonal antibody were generated in our laboratory. PDCoV N antibody (PDCoV11-M) was purchased from Alpha Diagnostic Intl. Inc, USA. Rabbit anti-Importin α1 (10819-1-AP), rabbit anti-STAT2 (16674-1-AP) were purchased from Proteintech Group, Inc., China. Rabbit anti-Importin α3 (A2026), rabbit anti-Importin α4 (A8347), rabbit anti-Importin α5 (A24117), rabbit anti-Importin α6 (A7331), rabbit anti-Importin α7 (A8347), rabbit anti-HA (AE105), rabbit anti-Myc (AE070), rabbit anti-β-Actin (AC026) were purchased from ABclonal Technology Co., Ltd., China. Rabbit anti-STAT1 (#14994) was purchased from Cell Signaling Technology, Inc., USA. Rabbit anti-IRF3 (ab68481) and rabbit anti-p65 (ab32536) were purchased from Abcam plc., UK. Mouse anti-Flag (M185-3L) was purchased from Medical & Biological Laboratories Co., Ltd., Japan. The dilution of antibodies is summarized in S1 Table. Goat anti-rabbit IgG (H+L) (AS014), and goat anti-mouse IgG (H+L) (AS003) conjugated with HRP were purchased from Jackson ImmunoResearch Laboratories Inc., USA. Alexa Fluor goat anti-rabbit-488 (A-11034), Alexa Fluor goat anti-rabbit-594 (A-11037), Alexa Fluor goat anti-mouse-488 (A-11029), and Alexa Fluor goat anti-mouse-594 (A-11005) were purchased from Invitrogen, USA. Poly(I:C) (31852-29-6) was purchased from Invitrogen, USA. Recombinant human IFN-β protein (#8499-IF) was purchased from Bio-Techne R&D Systems, USA. Recombinant Human TNF-α (P00029) was purchased from Beijing Solarbio Science & Technology Co.,Ltd., China.

### Plasmids construction

The sequences encoding Flag-tagged PEDV nsp1, nsp2, PLpro-TM, PLpro, nsp4, nsp5, nsp6, nsp7, nsp8, nsp9, nsp10, nsp12, nsp13, nsp14, nsp15, nsp16, S, E, M, N, and ORF3, were generated by amplifying cDNA from PEDV HLJBY strain using corresponding primers and cloning into PXJ40 with 2×MultiF Seamless Assembly Mix (RK21020, ABclonal, China), with Flagbtag at the N-terminus. The sequences encoding MHV PLpro-TM, TM-truncated MHV-PLpro, IBV PLpro-TM, TM-truncated IBV-PLpro, PDCoV PLpro-TM, and TM-truncated PDCoV-PLpro, were amplified from MHV, IBV Beaudette strain, or PDCoV cDNA templates using gene-specific primers, followed by cloning into the PXJ40 vector, with Flag tag at the N-terminus. The catalytic mutant plasmid of PEDV PLpro-TM (C99A, H258A and D271A), and IBV PLpro-TM (C101A, H264A and D275A), were cloned using the Mut Express II Fast Mutagenesis Kit V2 (C214, Vazyme, China). Sequence encoding Importin α1 and Importin α4 were amplified from HEK-293T cells cDNA and cloned into pCMV (with HA tag at the N-terminus and Myc-tag at the C-terminus). SV40-NLS-GFP were constructed by Dr. Xue Wenxiang [50]. The PY-NLS sequence (GGTAGTGGTTTTGGGAATTACAACAATCAGTCTTCAAATTTTGGACCCATGAAGGGAGGAAATTTTGGAGGCAGAAGCTCTGGCCCCTAT) were synthesized by Sangon Bioengineering Co., Ltd., Shanghai, China, and ligated into the pEGFP-N1 vector. The pCMV-Importin α1 G72A, G84A, G119A, G129A and G162A were mutated using the Mut Express II Fast Mutagenesis Kit V2. The primers used to generate the above plasmids are listed in S2 Table.

### Western blotting analysis

Cells were seeded in 6-well plates and infected with PEDV HLJBY strain, IBV Beaudette strain, PDCoV, or transfected with various plasmids, according to experimental requirements. At indicated h.p.i. or h.p.t., cells were harvested and lysed in 2 × protein loading buffer (20 mM Tris-HCl, 2 % SDS, 100 mM DTT, 20 % glycerol, 0.016 % bromophenol blue) and incubated in a 100 °C metal bath for 5 min to fully denature the proteins. The cell lysates were then subjected to centrifugation at 12000 rpm for 5 min to remove the cell debris. The supernatants were resolved on a 10% or 12% SDS-PAGE and transferred to a nitrocellulose membrane (Millipore, USA). Membranes were blocked in blocking buffer (5 % fat free milk, TBS, 0.5 % Tween 20) for 1 h, followed by overnight incubation at 4 °C with primary antibodies diluted in dilution buffer (WB500D, NCE, China) (as indicated in S1 table). After washing with TBST (0.5 % Tween 20 in TBS) for three times, the membranes were then incubated with secondary antibodies conjugated with HRP (Invitrogen, USA) diluted 1:10,000 in TBST for 1 h at room temperature, followed by washing three times with TBST. The antibody probed proteins were visualized using Tanon 4600 Chemiluminescent Imaging System (Bio Tanon, China). Image J program (NIH, USA) was used to quantify the intensities of corresponding bands on the Western blot according to the manufacturer’s instructions.

### Indirect immunofluorescence analysis

Cells were seeded onto chamber slides and transfected with corresponding plasmids, or infected with PEDV HLJBY or IBV Beaudette strain, according to experimental requirements. At the indicated time points, cells were transfected with poly(I:C) (20 μg/mL, 6 h), or treated with IFN-β (1000 IU/mL, 45 min) or TNF-α (20 ng/mL, 45 min). PBS treatment was used as the negative control. Following treatment, cells were fixed with 4 % paraformaldehyde for 15 min and permeabilized with 0.5 % Triton X-100 for 15 min at room temperature. After three times washes with PBS, cells were incubated in blocking buffer (3 % BSA in PBS) for 1 h at 37°C. Cells were then incubated with the primary antibody diluted in blocking buffer (as indicated in S1 Table) overnight at 4°C, followed by incubation with Alexa Fluor-conjugated secondary antibody diluted with 1:500 in blocking buffer for 1 h at 37°C. In case of double staining, cells were incubated with the other primary antibody, followed by incubation with the corresponding conjugated secondary antibody and incubated. Between and after each incubation step, the cell monolayer was washed three times with washing buffer. Cell nuclei were stained with DAPI Fluoromount-G™ (36308ES, Yease, China) for 15 min and the slides were mounted with mounted buffer. Finally, the subcellular localization of corresponding proteins was examined using a Zeiss LSM880 confocal microscope. Protein fluorescence intensity was quantified using Image J software (National Institutes of Health, USA). Statistical analysis of the nuclear/cytoplasmic fluorescence intensity ratio for the target protein was performed using GraphPad Prism 8 software. Quantitative results were from three fields of view within one representative single experiment, with error bars representing the SD of the nuclear/cytoplasmic fluorescence distribution histogram for the target protein. The significance of differences between two groups was evaluated using a two-tailed independent Student’s t-test. A *P-*value of less than 0.05 was considered statistically significant. Statistical significance levels are denoted as follows: *, *P* < 0.05; **, *P* < 0.01; ***, *P* < 0.001; ****, *P* < 0.0001.

### RNA extraction and real-time quantitative RT-PCR analysis

HEK-293T cells, PK-15 cells or DF-1 cells were seeded in 6-well plates and transfected with corresponding plasmids. At 24 h.p.t., cells were transfected with poly I:C (20 μg/mL, 8 h), or treated with IFN-β (1000 IU/mL, 8 h), or treated with TNF-α (20 ng/mL, 8 h). PBS treatment was included in parallel experiments as a negative control. Total cellular RNA was extracted using FreeZol Reagent (R711-01, Vazyme, China). Cell pellets or viral suspensions were lysed in 500 μL of FreeZol Reagent and incubated at room temperature for 5 min. Subsequently, 150 μL of Dilution Buffer was added, and the mixture was vigorously vortexed and allowed to stand for another 5 min. The samples were then centrifuged at 12,000 rpm for 15 min at room temperature. A 500 μL aliquot of the supernatant was transferred to a fresh tube, mixed with an equal volume of isopropanol by inversion, and incubated at room temperature for 30 min. After centrifugation at 12,000 rpm for 15 min at room temperature, the supernatant was carefully discarded, and the RNA pellet was washed twice with 1 mL of 75% ethanol. To remove residual ethanol, the tube was briefly centrifuged for 2 min at room temperature, and the remaining liquid was aspirated with a pipette tip. The pellet was air-dried, dissolved in 20 μL of RNase-free ddH₂O, and allowed to stand for 2 min. The resulting total RNA was stored at –80 °C until further use.

By using above RNA as template, the cDNA was synthesized by reverse transcription using the EasyScript One-Step gDNA Removal and cDNA Synthesis SuperMix kit (AE311, Trans, China) with oligo dT primer. The cDNA was then served as a template for real-time qPCR using SYBR green master mix (SB-Q204, Share-bio, China) and corresponding gene specific primers (IFN-β, ISG56 or IFITM3, and IL-8). Real-time qRT-PCR was conducted in the CFX-96 Bio-Rad instrument (Bio-Rad, USA) and the specificity of the amplified PCR products was confirmed by melting curve analysis after each reaction. The primers used to detect antiviral genes (IFN-β, ISG56 or IFITM3, and IL-8) are listed in S3 Table. Statistical analysis was performed using GraphPad Prism 8 software. The data are presented as bar graphs, with error bars representing the SD of three technical replicates within single representative experiment. Significance was determined using the two-tailed independent Student’s t-test (*P* < 0.05) between two groups. Statistical significance was denoted as follows: *, *P* < 0.05; **, *P* < 0.01; ***, *P* < 0.001; ****, *P* < 0.0001.

### Co-immunoprecipitation (Co-IP)

HEK-293T cells or Vero cells cultured in 6 cm or 10 cm plates were transfected with corresponding plasmid for 24 h or infected with IBV for 18 h. Cells were lysed using RIPA Lysis Buffer (P0013D, Beyotime, China) supplemented with 1 mM phenylmethylsulfonyl fluoride (PMSF) (ST506, Beyotime, China). The cell lysates were centrifuged at 12,000 rpm for 15 min, and the supernatant were incubated with 2 μg of anti-Flag monoclonal antibody (mouse), or 4 μg of IBV nsp3 polyclonal antibody (mouse) conjugated with Dynabeads Protein G Magnetic Beads (10004D, Invitrogen, USA), or anti-HA magnetic Beads (SB-PR003, Share-bio, China), for 2 h at room temperature with gentle rotation. After incubation, the beads were washed six times (5 min each time) with RIPA buffer and then precipitated using a magnetic stand. The beads were resuspended in 40 μL of RIPA lysis buffer and 5 × SDS loading buffer (P0015L, Beyotime, China), followed by boiling at 100 °C for 5 min. Following 12000 rpm centrifugation for 5 min, the supernatants were subjected to Western blot analysis.

## Supporting information

**S1 Table. Dilution of primary antibodies.**

**S2 Table. Primer sequences used for plasmid construction.**

**S3 Table. Primer sequences used for real-time qPCR.**

## Acknowledgments

We gratefully acknowledge Prof. Dingxiang Liu (South China Agricultural University, China) for providing the IBV Beaudette strain. We are also thankful to Assoc. Prof. Changchao Huan’s laboratory (Yangzhou University, China) for supplying the PEDV HLJBY strain, and to Prof. Tongling Shan, Prof. Yanjun Zhou, and Prof. Fei Gao (all from Shanghai Academy of Agricultural Sciences, CAAS, China) for providing the PDCoV strain, the PEDV anti-N protein antibody, and the LLC-PK1 cells, respectively. We also thank Dr. Huan Wang, Dr. Bo Gao, and Dr. Xiaoqian Gong (Shanghai Academy of Agricultural Sciences, CAAS, China) for constructing the Flag-tagged IBV protein-encoding plasmids. Finally, we appreciate the technical assistance of Hongyang Liu and Qianxing Hou (Shanghai Academy of Agricultural Sciences, CAAS, China) in cell culture and sample collection.

## Author Contributions

**Conceptualization:** Jiehuang Wang, Ying Liao.

**Formal analysis:** Jiehuang Wang, Ying Liao.

**Funding acquisition:** Ying Liao.

**Investigation:** Jiehuang Wang, Wenxiang Xue.

**Project administration:** Ying Liao.

**Resources:** Yingjie Sun, Xusheng Qiu, Cuiping Song, Lei Tan.

**Supervision:** Ying Liao, Yingjie Sun, Chan Ding.

**Writing – original draft:** Jiehuang Wang, Ying Liao.

**Writing – review & editing:** Ying Liao.

